# Cellular and molecular mechanisms of photoreceptor tuning for prey capture in larval zebrafish

**DOI:** 10.1101/744615

**Authors:** Takeshi Yoshimatsu, Cornelius Schröder, Noora E Nevala, Philipp Berens, Tom Baden

## Abstract

In the eye, the function of same-type photoreceptors must be regionally adjusted to process a highly asymmetrical natural visual world. Here we show that UV-cones in the larval zebrafish *area temporalis* are specifically tuned for UV-bright prey capture in their upper frontal visual field, which uses the signal from a single cone at a time. For this, UV-detection efficiency is regionally boosted 42-fold. Next, *in vivo* 2-photon imaging, transcriptomics and computational modelling reveal that these cones use an elevated baseline of synaptic calcium to facilitate the encoding of bright objects, which in turn results from expressional tuning of phototransduction genes. Finally, this signal is further accentuated at the level of glutamate release driving retinal networks. These regional differences tally with variations between peripheral and foveal cones in primates and hint at a common mechanistic origin. Together, our results highlight a rich mechanistic toolkit for the tuning of neurons.

## INTRODUCTION

In vision, photoreceptors drive the retinal network through continuous modulations in synaptic release (Baden et al., 2013a; Heidelberger et al., 2005; Lagnado and Schmitz, 2015; Moser et al., 2019; Regus-Leidig and Brandstätter, 2012; Thoreson, 2007). However, how changes in incoming photon flux lead to changes in the rate of vesicle fusion at the synapse varies dramatically between photoreceptor designs (Bellono et al., 2018; Sterling and Matthews, 2005; Thoreson, 2007). For example, in the vertebrate retina, the slow rod-photoreceptors typically have large outer segments and high-gain intracellular signalling cascades to deliver single-photon sensitivity critical for vision at low light (Field et al., 2005; Lamb, 2016; Yau and Hardie, 2009). In contrast, cone-photoreceptors are faster, have smaller outer segments and lower-gain cascades to take over where rods saturate (Lamb, 2016; Yau and Hardie, 2009). Clearly, matching the properties of a given photoreceptor type to a specific set of sensory tasks critically underpins vision. However, these visual requirements can differ dramatically across the retinal surface and the corresponding position in visual space (Baden et al., 2013b; Hardie, 1984; Land and Nilsson, 2012; Sancer et al., 2019; Yilmaz and Meister, 2013; Zimmermann et al., 2018). For efficient sampling (Cronin et al., 2014; Land and Nilsson, 2012), even cones of a single type must therefore be functionally tuned depending on their retinal location.

Indeed, photoreceptor tuning – even within-type – is a fundamental property of vision in both invertebrates (Hardie, 1984; Sancer et al., 2019) and vertebrates (Baden et al., 2013b; Baudin et al., 2019; Sinha et al., 2017). Even primates make use of this trick: foveal cones have longer integration times than their peripheral counterparts, likely to boost their signal to noise ratio as in the foveal centre retinal ganglion cells do not spatially pool their inputs (Baudin et al., 2019; Sinha et al., 2017). Understanding the mechanisms that underlie such functional tuning will be important for understanding how sensory systems can operate in the natural sensory world, and how they might have evolved to suit new sensory-ecological niches (Cronin et al., 2014; Lamb et al., 2007; Land and Nilsson, 2012; Yau and Hardie, 2009).

Here, we show that UV-cones in the *area temporalis* (Schmitt and Dowling, 1999) (“strike zone, SZ” (Zimmermann et al., 2018)) of larval zebrafish are selectively tuned to detect microorganisms that these animals feed on (e.g. paramecia) (M, 2000; Spence et al., 2008). The performance of this prey-detection system is remarkable: Our calculations show that the sequential signals from single UV-cones at a time, each activated for mere tens of milliseconds, must suffice to trigger prey-orientation behaviour. Zebrafish achieve this performance through a series of region-specific photoreceptor optimisations. Using *in-vivo* 2-photon calcium and glutamate imaging we show that UV-cones only in the strike zone combine a grossly enlarged outer segment, an elevated synaptic calcium baseline and a slowed response recovery from light-activation to maximise the detectability of rapidly moving UV-bright paramecia (Jung et al., 2014). In contrast, UV-cones in other parts of the eye preferentially respond to the sudden absence of light, likely to support a balance of predator detection and colour vision (Cronin and Bok, 2016; Zimmermann et al., 2018). Using transcriptomics, pharmacological manipulation and computational modelling we go on to identify the cellular, molecular and synaptic mechanisms that underlie these specialisations.

## RESULTS

### Larval zebrafish prey capture must use UV-vision

Larval zebrafish prey capture is elicited by a bright spot of light (Bianco et al., 2011; Semmelhack et al., 2014), in line with the natural appearance of their prey items (e.g. paramecia) in the upper water column of shallow water when illuminated by the sun (Zimmermann et al., 2018) (Fig. 1a). To the human observer with comparatively long-wavelength vision (Nathans, 1999) these organisms are largely transparent when viewed against a back-light (Johnsen and Widder, 2001). However, previous work suggests that zooplankton like paramecia scatter light in the UV-band (320-390 nm) and thus appear as UV-bright spots (Novales Flamarique, 2012, 2016; Zimmermann et al., 2018).

**Figure 1.**
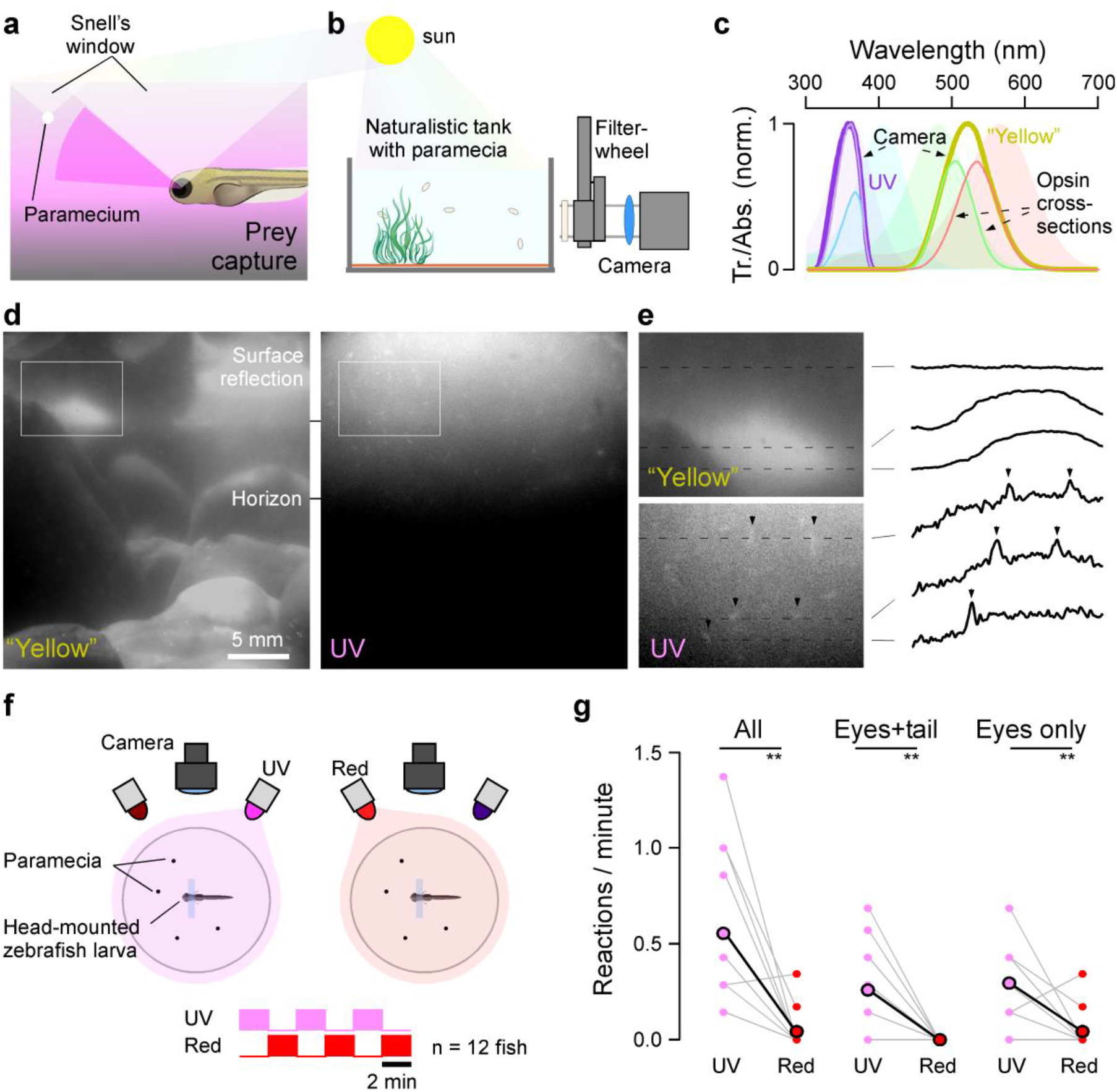
UV-light greatly facilitates visually guided prey capture in larval zebrafish. **a**, Schematic representation of visual prey capture by larval zebrafish. **b**, Set-up for filming paramecia. A filter wheel equipped with UV and yellow bandpass filters was positioned in front of the CCD camera to image paramecia in a naturalistic tank in the sun. **c**, Peak-normalized spectra for the UV and “yellow” channels (thick lines, Methods) superimposed on the zebrafish’ four opsin absorption spectra (shadings). The spectral overlap between the UV and “yellow” channel with each opsin is indicated (thin lines). “Abs.” = absorption, “Tr.” = transmittance. **d**, Example frames from the “yellow” and UV channels taken consecutively from the same position. **e**, Zoom in from (d), with line profiles extracted as indicated. Arrowheads highlight paramecia visible in the UV channel. See also Vid. S1. **f**, Schematic of behavioural setup: individual larval zebrafish (7-8 dpf) in the presence of free-swimming paramecia were head-mounted and filmed from above, with infrared illumination from below. In addition, top-illumination was provided by either UV (374±15 nm) or red (620±10 nm) LEDs as indicated. g, Zebrafish consistently responded more readily to passing paramecia with either full prey-capture bouts (eye convergence + tail flicks) or with tracking movements during UV-illumination periods. Paired t-test, n=12: p=0.0014, 0.0039, 0.0025 as indicated, respectively.

To explicitly test this idea, we custom-built a camera system with a UV and a “yellow” channel aligned with the zebrafish UV- and red/green opsin absorption spectra, respectively (Chinen et al., 2003). We used this system to film free-swimming paramecia in a naturalistic tank placed outdoors under the mid-day sun (Fig. 1b-e, Vid. S1, Methods). While the “yellow” image provided good spatial detail of the scene’s background and surface water movements, paramecia were difficult to detect amongst the background clutter (Fig. 1d, left). In contrast, the UV-channel was dominated by a vertical brightness gradient of scattered light which almost completely masked the background. Superimposed on this gradient, the upper water column readily highlighted individual paramecia as bright moving spots (Fig. 1d right, 1e). In agreement, zebrafish use their upper-frontal visual field to detect and capture prey (Bianco et al., 2011; Mearns et al., 2019; Patterson et al., 2013), and inner retinal circuits that process this part of visual space exhibit a strong, regionally specific bias for UV-bright contrasts (Zimmermann et al., 2018). This confirmed that vastly different, and largely non-overlapping types of information (Cronin and Bok, 2016) are obtainable from these two wavebands available to the zebrafish larvae.

Notably, any differences between the UV and “yellow” waveband will be further exacerbated by the fish’s self-movements relative to the scene. These would add major brightness transitions in the “yellow”, but not the UV channel. Accordingly, under natural - rather than laboratory-controlled - viewing conditions paramecia are likely nigh-impossible to detect in the “yellow” waveband, but readily stand out in the UV. This strongly suggests that larval zebrafish must capitalise on UV-vision – rather than achromatic or long-wavelength vision - to support visual prey-detection in nature (Cronin and Bok, 2016; Novales Flamarique, 2016; Zimmermann et al., 2018).

Indeed, UV-illumination strongly facilitated behavioural performance: Head-mounted 7-8 days post fertilisation (*dpf*) larvae in the presence of free-swimming paramecia exhibited significantly more prey-capture attempts when illuminated with UV-light (374 nm), compared to red light (620 nm) (Fig. 1f,g).

### Single UV-cones signal the presence of prey

The ~300 μm diameter eyes of larval zebrafish necessarily offer limited spatial resolution (Haug et al., 2010), meaning that visually detecting their even smaller prey presents a substantial challenge. We therefore set out to determine the maximal numbers of UV-cones the fish can use for this task. At 8 *dpf*, larval zebrafish have ~2,400 UV-cones distributed per eye. These are unevenly distributed and exhibit a three-fold elevation in the centre of the strike zone (SZ) (Zimmermann et al., 2018) which in visual space is located at 38° azimuth and 27° elevation relative to the centre of the monocular field (Fig. S1a). At rest and during hunting, larval zebrafish converge their 169°±4.9° (n=4) field of view eyes by ~36° and ~76° (Bianco et al., 2011; Patterson et al., 2013; Trivedi and Bollmann, 2013) to afford a frontal binocular overlap of 26° and 66°, respectively (Fig. S1b-d). Based on these numbers, we computed the spatial detection limits of the UV-detector array across both eyes (Fig. 2a-d, Fig. S1 a-d).

**Figure 2.**
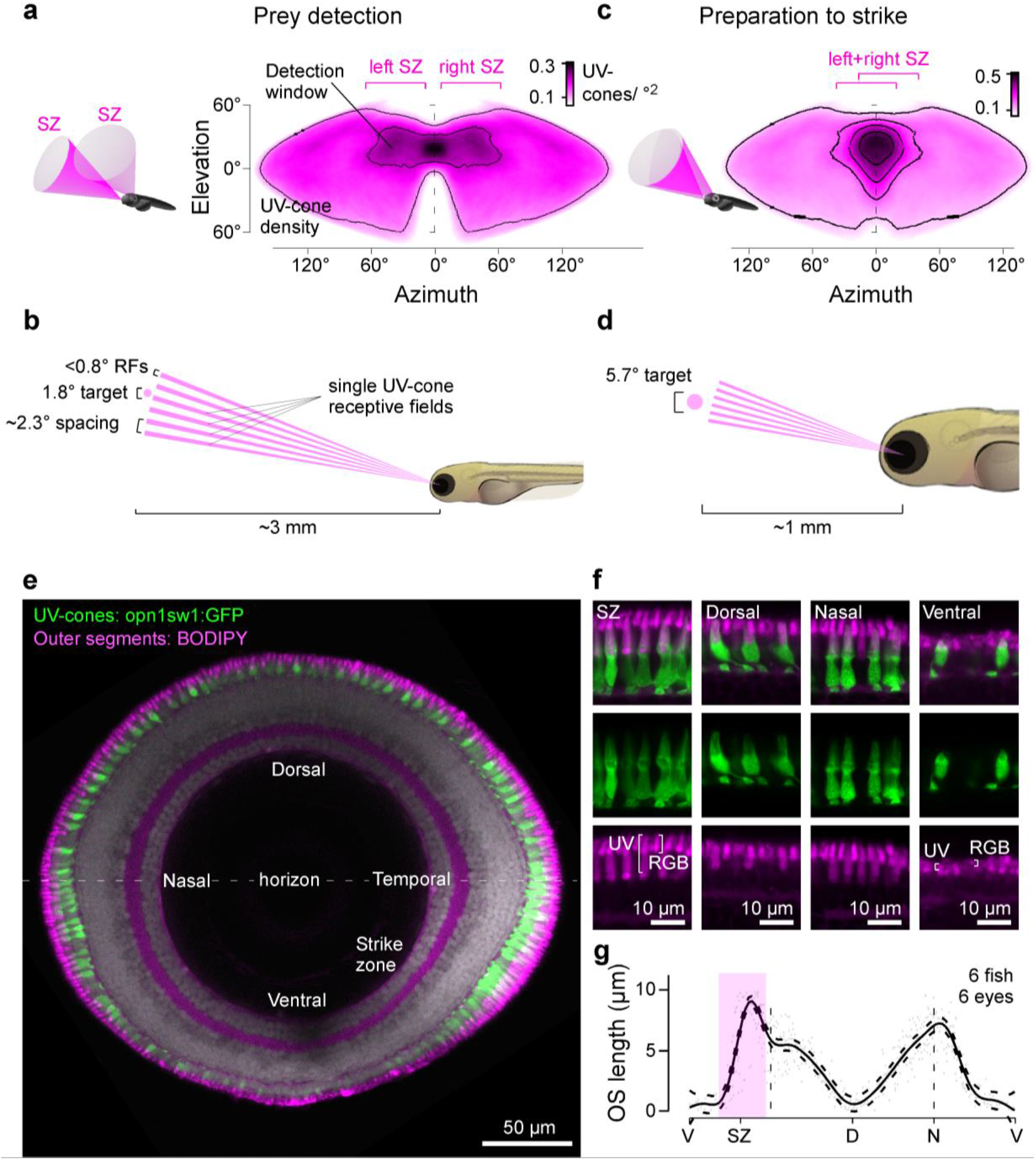
The detector hardware for UV vision in larval zebrafish. **a-d**, UV-cone density projected into sinusoidal map of visual space when eyes are in resting position for initial prey detection (a) and once converged for prey localisation following detection (c). A 100 μm paramecium is too small for reliable detection at ~3 mm distance and can therefore only be seen by a single UV-cone at a time (b). Even at ~1 mm strike-distance it covers at most a handful of UV-cones per eye (d). 3D-Schematics (a,c) illustrate approximate visual space surveyed by the two SZs. Scale bars in UV-cones / °. See also Fig. S1a-d. **e**, Sagittal section across the eye with outer segments (OS) stained by BODIPY (magenta) and UV-cones expressing GFP (green, Tg(opn1sw1:GFP)) in an 8 dpf larva. **f**, Higher magnification sections from (e). Note that BODIPY stains the OS’s of all photoreceptors, as well as the spot-like pocket of mitochondria immediately below the OS (Fig. S1e-g). Note also that region-specific OS enlargements are restricted to UV cones. g, Mean and 95% confidence intervals of UV-cone OS lengths across the eye. V, ventral; SZ, strike zone; D, dorsal; N, nasal. 3D fish model by MY Zimmermann.

Before initiating the actual strike, and prior to converging their eyes, zebrafish must first detect their prey (Gahtan, 2005; McElligott and O’Malley, 2005). This mostly occurs within the upper visual field (~30° elevation (Mearns et al., 2019)), where the UV-signal from paramecia is particularly prominent (Fig. 1d,e). Within this region, prey-detection performance is highest when the target is laterally displaced from the centre of the binocular visual field by ~23° (Mearns et al., 2019). This same region was surveyed by each eye’s SZ (Fig. 2a, Fig. S1a,b), confirming that zebrafish indeed capitalise on the elevated UV-cone density in this part of the eye for prey detection. However, with a mean SZ UV-cone spacing of 0.19 cones/°^2^, and a UV-cone receptive field diameter of ~0.76°, even at its peak this UV-detector array nevertheless dramatically undersamples visual space for this critical behavioural task (Fig. 2b): Larval zebrafish can detect <100 μm prey (Lawrence, 2007; Wilson, 2012) at up to 3.25 mm (Bianco et al., 2011) distance, where it subtends a visual angle of only 1.8°. This is more than two times smaller than required for reliable detection at the Nyquist limit. It therefore follows that zebrafish cannot use more than a single UV-cone at a time to trigger the initial behavioural response.

Once this prey is detected, zebrafish orient towards it and converge their eyes (Bianco et al., 2011; Gahtan, 2005; Jouary et al., 2016; McElligott and O’Malley, 2005; Mearns et al., 2019; Patterson et al., 2013; Trivedi and Bollmann, 2013). This brings both SZs into near perfect alignment directly in front of the fish, thus enabling stereoptic estimation of exact prey position for subsequent capture (Patterson et al., 2013) (Fig. 2c, Fig S1c,d). The actual strike is then initiated at a distance of ~1 mm (Patterson et al., 2013), when a 100 μm paramecium subtends a visual angle of ~5.7° (Fig. 2d). At this angular size, it reliably covers 2-3 UV-cones per eye, yet rarely substantially more. Taken together, single UV-cones in the SZ therefore likely underlies initial prey detection triggering prey-orientation behaviour, and even subsequent distance estimation leading to triggering the actual strike can only be supported by a handful of UV-cones per eye.

### UV-cone outer segment size varies more than ten-fold across the eye

As single cones need to suffice for prey detection, and in view of the relatively low UV-signal in natural light (Losey et al., 1999; Zimmermann et al., 2018), UV-cones in the strike zone must be able to absorb photons with high efficiency to support hunting behaviour. In contrast, UV-cones outside the strike zone might be able to afford lower efficiency and thus conserve space and energy as it is possible to pool the coincident signals from multiple UV-cones, for example for UV-dark silhouette based predator detection (Cronin and Bok, 2016). A simple way to increase a vertebrate photoreceptor’s photon catch efficiency is to enlarge its outer segment which houses the phototransduction machinery (de Busserolles et al., 2014; Warrant and Nilsson, 1998). To test this, we genetically labelled all UV-cones (green), stained outer segments of all cones using the membrane dye BODIPY (magenta) and assessed their morphology using confocal imaging (Fig. 2e-g). This revealed more than ten-fold variations in outer segment lengths. SZ UV-cones had the longest outer segments (9.0±0.4 μm) while the immediately neighbouring ventral UV-cones had the shortest (0.6±0.8 μm). A secondary peak occurred in nasal UV-cones (7.0±0.5 μm) which survey the outward horizon – possibly to also support the UV-driven chromatic circuits in this part of the eye (Zimmermann et al., 2018). In cyprinid photoreceptors, photon catch efficiency (*F*) scales as a function of outer segment length (*l*) as

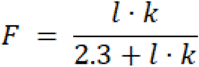

where *k* is the photoreceptor-type specific absorption coefficient of 0.03^-μm^ (Warrant and Nilsson, 1998). Accordingly, the observed variation in outer segment length from 0.6 to 9.0 μm should lead to a ~14-fold boost in photon catch efficiency for SZ cones. Together, the combination of UV-cone density across the eye (factor 3), outer segment length (factor 14) and binocular superposition of the two eyes’ SZs during hunting (factor 2) should therefore lead to a 42 (not-converged) to 84-fold (converged) variation in UV-sensitivity across the visual field.

Finally, located just beneath each outer segment, SZ UV-cones also had consistently enlarged ellipsoid bodies (Fig. 2f, Fig. S1e-g). These structures house the mitochondria that power phototransduction (Giarmarco et al., 2017; Okawa et al., 2008), but they might further act as micro-lenses to focus additional light onto outer segments (Knabe et al., 1997). With a more than five-fold variation in ellipsoid body 2D-area across the eye (Fig. S1g), any such focussing effect would further boost UV-detection capacity of the SZ. We next asked how these anatomical differences might be reflected at the level of UV-light responses across the *in vivo* eye.

### SZ UV-cones are light-biased and have long integration times

To measure UV-cone light responses *in vivo*, we expressed the synaptically tagged fluorescent calcium biosensor SyGCaMP6f (Dreosti et al., 2011) in all UV-cone pedicles. We co-expressed mCherry (Shaner et al., 2004) under the same opn1sw1 promotor (Takechi et al., 2003) without synaptic tagging to reveal each cone’s full morphology and to confirm that SyGCaMP6f expression was restricted to the pedicles (Fig. 3a). 7-8 *dpf* larvae were imaged under 2-photon at 64×16 pixel resolution (62.5 Hz), capturing 1-5 UV-cone pedicles at a time. This allowed imaging light-driven cone-pedicle calcium in any part of the *in vivo* eye.

We first presented light- and dark- flashes from a constant UV-background (Methods). Prey capture behaviour can be initiated by the presentation of a bright spot as small as 2°, moving at a speed of 90 °/s (Semmelhack et al., 2014). Such a moving stimulus activates a single UV-cone for at most 30 ms if perfectly centred. At times, paramecia will however move somewhat slower (cf. Vid. S1), meaning that also slightly longer stimulus durations are meaningful for prey-detection. Accordingly, we presented light- and dark-flashes at varying durations. In an example recording, we observed that a SZ UV-cone indeed responded to 20 ms and 50 ms UV-light flashes, while a dorsal cone failed to exhibit a detectable response (Fig. 3b, Vid. S2). However, compared to the SZ UV-cone, the dorsal UV-cone responded much more strongly to a 200 ms dark flash.

**Figure 3.**
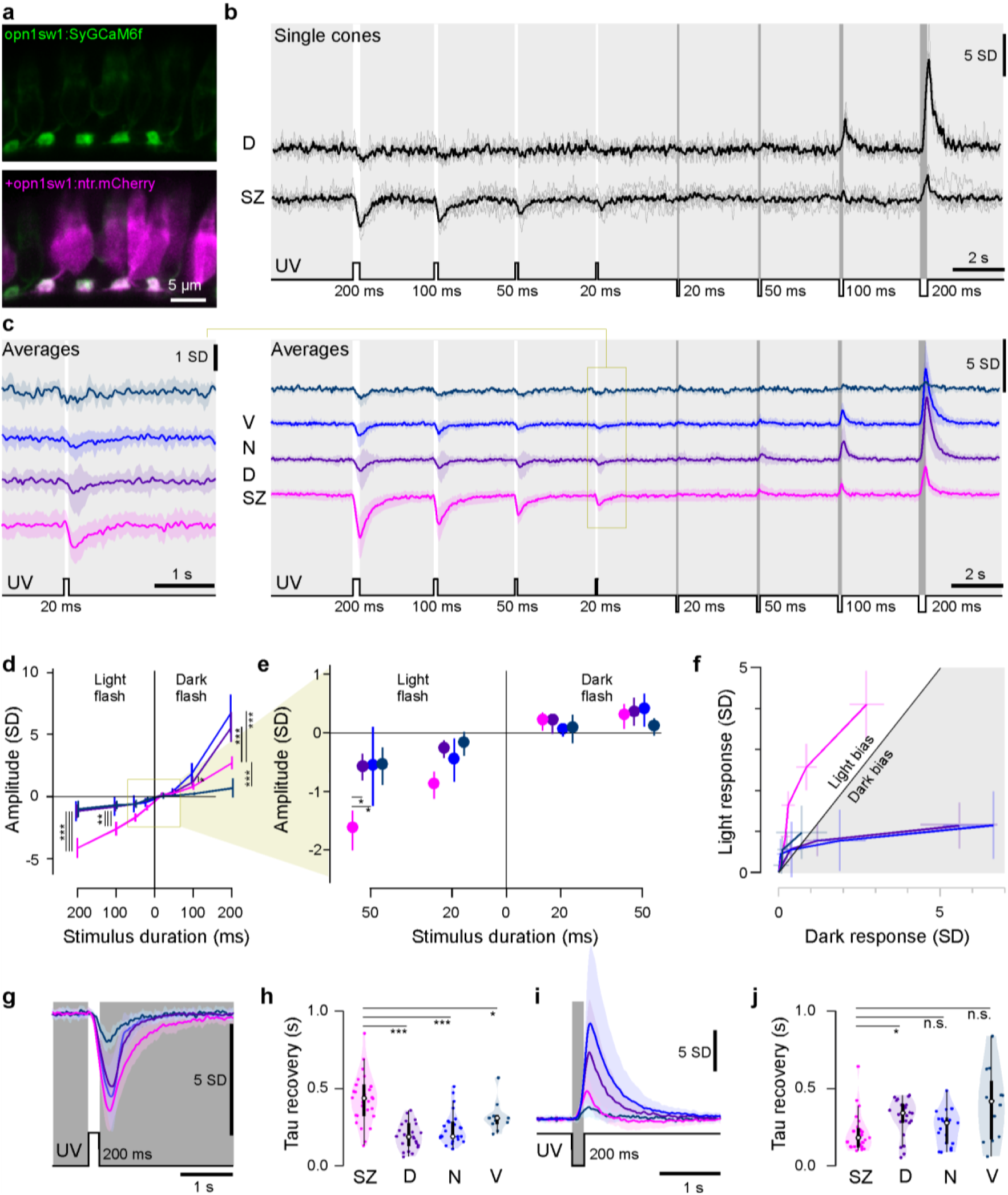
Imaging cone-calcium in the live eye. **a**, Confocal images of synaptically targeted GCaMP6f (green, Tg(opn1sw1:SyGCaMP6f)) in UV-cones (magenta, Tg(opn1sw1:nfsBmCherry)). **b**, Mean and single trial dorsal and SZ single cone 2-photon calcium responses to varying duration light- (6 x 10^5^ photon/s/μm^2^) and dark-steps (0 photon/s/μm^2^) from a constant UV background (2.4 x 10^4^ photon/s/μm^2^). **c**, Mean calcium responses to the same stimulus as in (b) from ventral, nasal, dorsal and strike zone cones (V, N, D, SZ; n = 9, 21, 23, 29, respectively). Shadings ±1 s.d.. Left shows an enlargement of the response to the 20 ms light step. **d**, Mean and 95% confidence intervals of peak amplitudes from (c). **e**, Enlargement from (d). All responses except nasal and ventral 20 ms dark-flash conditions were significantly different from zero (Mann-Whitney U-test). Within-condition pairwise comparisons across for SZ versus the other three zones are indicated with *, ** and *** (p<0.05, 0.01 and 0.001, respectively). P value adjustment: tukey method for comparing a family of 4 estimates. **f**, Light and dark responses from (c,d) plotted against each other for equivalent stimulus durations, with 95% confidence intervals indicated. **g**, Mean ±1 s.d. responses to a 200 ms flash of light (6 x 10^5^ photon/s/μm^2^) from darkness (0 photon/s/μm^2^). **h**, Box and violin plots of recovery time constants from (g). n = 29, 29, 23, 13 for SZ, D, N and V, respectively. **i, j**, as (g,h), but for an equivalent contrast dark-flash. n = 27, 24, 19, 13 for SZ, D, N and V, respectively. Mann-Whitney U-test *: p<0.02, ***: p<0.0001 (h,j). n.s.: not significant.

Across multiple such recordings, SZ cones consistently responded strongly to light flashes (Fig. 3c-f) including to the 20 ms condition (Fig. 3c,e), suggesting that SZ UV-cones are indeed well suited to detect the presence of UV-bright prey. In contrast, dorsal and nasal cones were dark biased (Fig. 3d,f) as would be useful to signal the presence of a UV-dark predator.

In addition to their unique light-bias, SZ UV-cones were also particularly slow to recover back to baseline following a light-flash (Fig. 3g,h). This prolonged response might aid temporal signal integration across multiple SZ UV-cones by postsynaptic circuits as the image of prey traverses the photoreceptor array. In contrast, recovery times from dark-flash responses were either similar or even slightly faster compared to the rest of the eye (Fig. 3i,j).

### UV-dependent prey detection is difficult outside the strike zone

Combining our data from the UV-cone distributions and *in-vivo* response properties we set-up a simple linear model to estimate how different types of UV-stimuli can be detected by the larval zebrafish’s monocular UV-detector array (Methods). For this, we first recorded the position of every UV-cone in a single eye and projected their 0.76° receptive fields into visual space (Fig. 4a, cf. Fig. 2a,c, Fig. S1a,c). We next computed a series of random-walk stimulus paths across this array by an assumed bright 2° target moving at an average speed of 100°/s and with approximately naturalistic turning behaviour (Jung et al., 2014; Shourav and Kim, 2017). This simulation confirmed our previous calculation that a single such target almost never (<0.1% of the time) covers two UV-cones at a time (Fig. 4b). In fact, most of the time (>60%), it covers zero UV-cones as it slips through gaps in the detector array. Even when adding all non-UV-cones (Methods), the maximal number of cones of any type covered at a time was three, with a single cone being the most likely incidence (~40%, Fig. 4b bottom).

**Figure 4.**
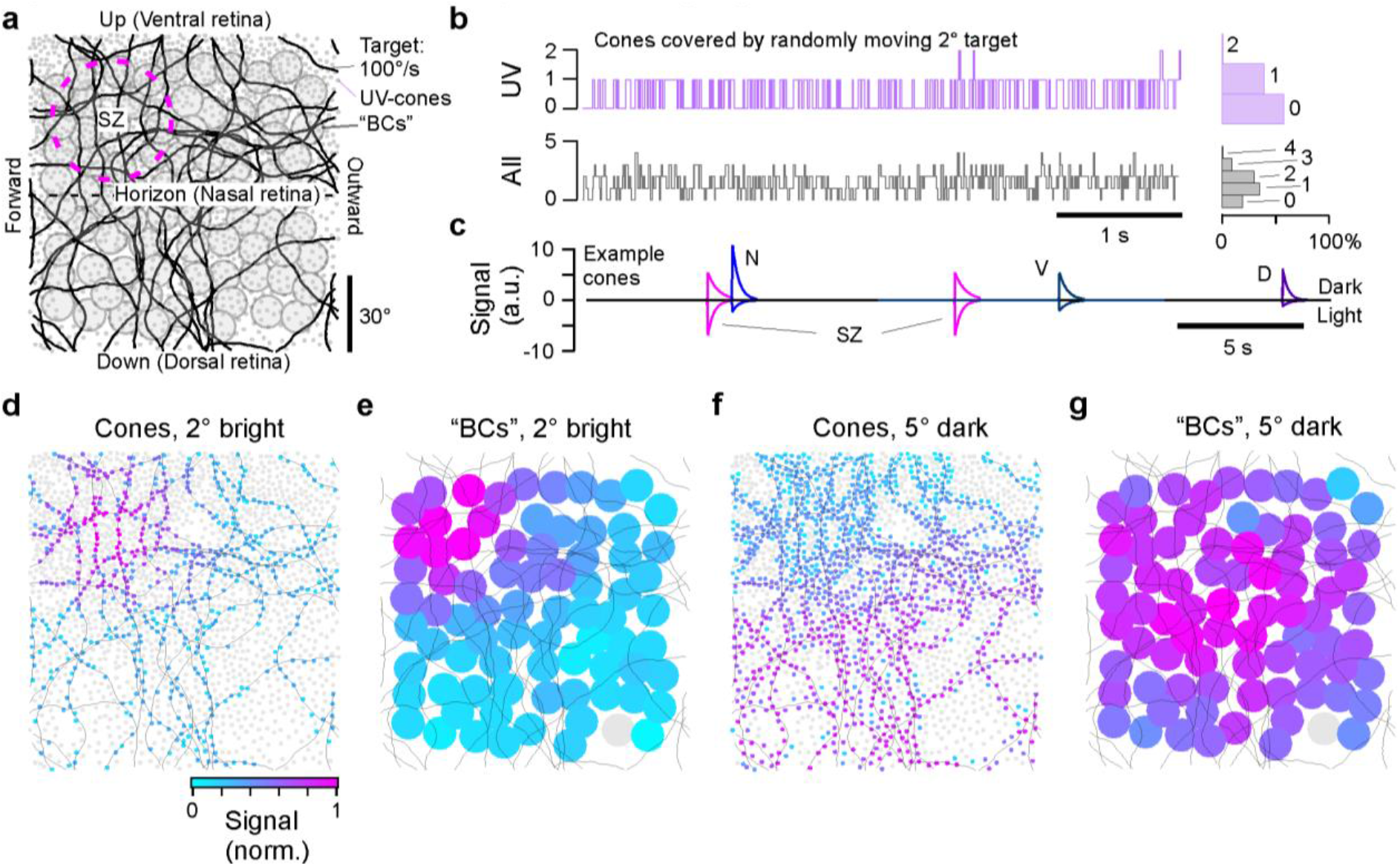
A model of UV-cone activation by a small moving target. **a**, Model set-up. Monocular UV-cone distribution across the visual field (grey dots) with model bipolar cell (BC) array superimposed (filled circles) and target paths black line. The strike zone was centred in the upper left quadrant, corresponding to the upper frontal visual field. **b**, Number of cones touched by a moving 2° plotted as an exeat over time, with histogram to the right. Top: UV-cones, Bottom: any-type cone. **c**, Time exeat of model cone activation of four example cones, taken from representative regions across the array. Responses below and above zero correspond to activation in response to a 2° bright and 5° dark target, respectively. **d**, Maximal activation levels of each cone over the full path for a 2° bright target, normalised to peak activation across the entire array. e, Activation of BCs driven by UV-cones in (d). **f,g**, as (d,e) but for a 5° dark target.

We then assigned response amplitudes and decay time-constants for both light- and dark-flashes based on our calcium imaging results to each UV-cone receptive field based on their position in the eye (Fig. 4c, cf. Fig. 3, Methods). For this, we also computed how an identically moving but larger (5°) dark target, meant to mimic a small or distant predator, activated UV-cones. In the model responses shown, single example cones from different parts of the retina responded sparsely to either object as it traversed their receptive fields. The model clearly predicted that the light object would be most detectable in the strike zone (Fig. 4d, Vid. S3). Adding even small amounts of noise would rapidly make all but SZ-based UV-light detection of this nature impossible. Any detectability difference would be further enhanced by a population of postsynaptic bipolar cells, here modelled to simply sum the signals from all UV-cones within a fixed radius. By integrating across more than one UV-cone, BCs also capitalise on the slower light recovery times of SZ UV-cones (Fig. 4e, cf. Fig. 3h, Methods). In contrast, the large dark object moving along the same path was detectable across the entire array (Fig. 4f). Here, the somewhat larger response amplitudes of dorsal UV-cones were approximately compensated for by the relatively greater number of UV-cones in the ventral half of the retina. This yielded an approximately homogeneous dark-response at the level of BCs across the entire visual field (Fig. 4g).

Taken together, the combination of differences in UV-cone density (Fig. 2a), outer segment size (Fig. 2e-g) and *in vivo* response properties at the level of presynaptic calcium driving release (Fig. 3) therefore strongly suggest that detection of paramecia using the UV-detector array will be strongly and specifically facilitated in the strike zone, and perhaps all but impossible in most other parts of the visual field.

We next explored the mechanisms underlying the dramatic shift in response-preference towards light stimuli by SZ UV-cones. For this, we returned to *in vivo* recordings of light-driven calcium across the eye.

### Differences in calcium baseline drive differential light-dark responses

To simultaneously record from all ~120 UV-cone pedicles in the sagittal plane at single-synapse resolution, we turned to higher spatial resolution scans of the full eye (Methods). In this configuration, the basal brightness of the SyGCaMP6f signal under a constant UV-background was consistently elevated in the strike zone (Fig. 5a). This brightness gradient was not related to differential SyGCaMP6f expression levels: When the same animal was fixed following live imaging and stained against the GFP fraction of SyGCaMP6f, the regional brightness differences were gone (Fig. 5b). This suggests that the SyGCaMP6f signal elevations in the live eye were linked to constitutive variations in UV-cone pedicle calcium baseline (Fig. 5c). We therefore further explored how calcium baseline varies between UV-cones, and how this in turn might affect their ability to encode light- and dark-stimuli.

**Figure 5.**
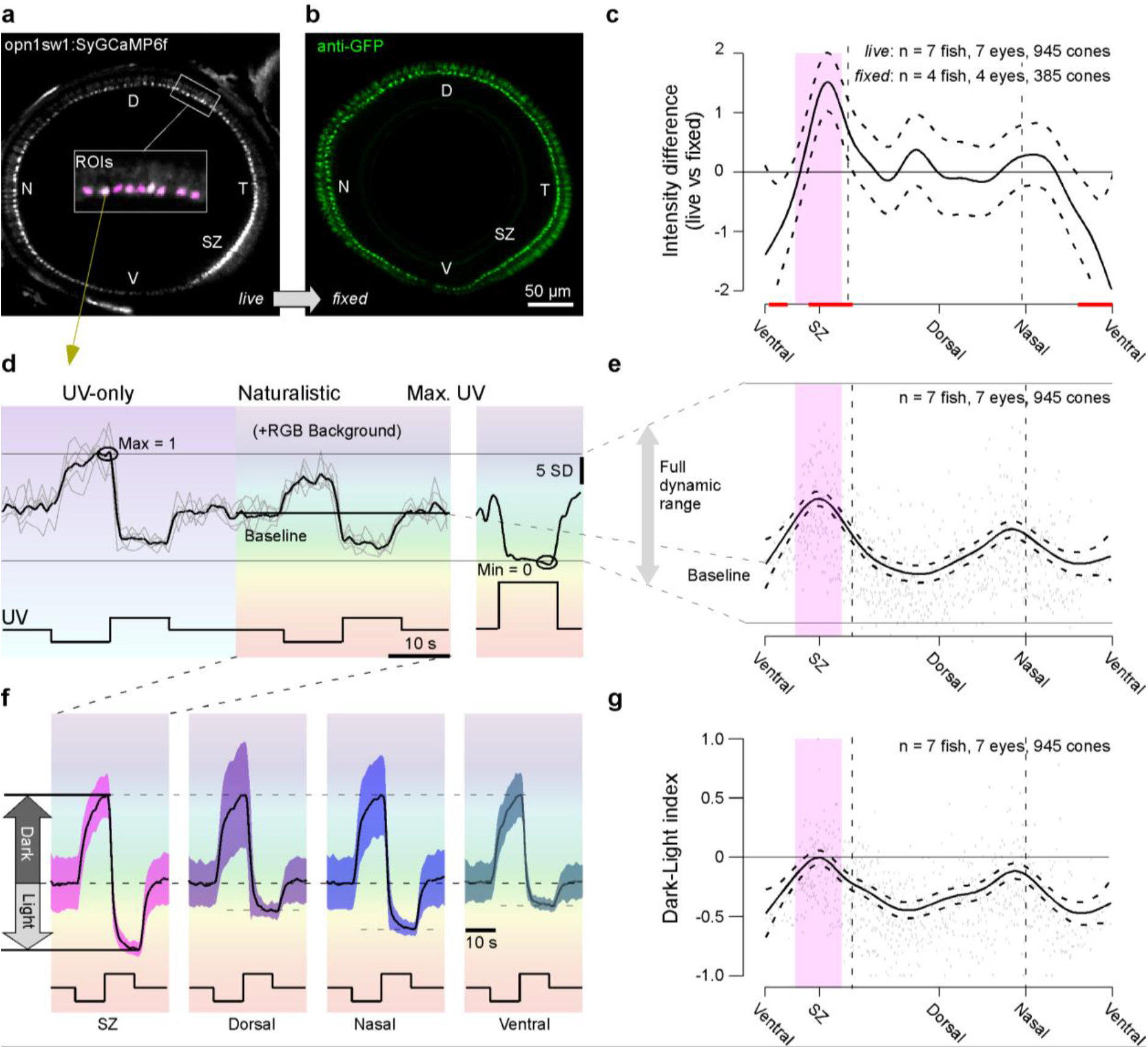
Calcium baseline predicts dark-light responses. **a,b**, Whole eye sagittal view of UV-cone SyGCaMP6f in live Tg(opn1sw1:SyGCaMP6f) zebrafish under 3 x 10^5^ photon/s/μm^2^ UV background light (a) and after immunostaining against SyGCaMP6f using anti-GFP antibody (b). **c**, Mean and 95% confidence intervals of the difference between live SyGCaMP6f signal per cone as in (a) and fixed signal as in (b), with red lines indicating regions that were significantly different from zero. **d**, Example mean and individual trial single cone response to 0 photon/s/μm^2^ dark- and 6 x 10^5^ photon/s/μm^2^ light-steps from a constant brightness UV 3 x 10^5^ photon/s/μm^2^ without and with spectrally broad background light. After five repeats, a 1.5 x 10^7^ photon/s/μm^2^ UV light step was presented to drive calcium to a minimum (right). **e**, Mean and 95% confidence interval of calcium baseline relative to the full dynamic range as indicated, with single datapoints in the back. **f**, Mean ± 1 s.d. calcium responses to light- and dark-contrasts with naturalistic RGB background light across all UV-cones in specified regions. Traces were shifted and scaled to align the baseline and peak dark-response. **g**, Mean and 95% confidence intervals of dark-light index (DLi) with single datapoints in the back.

To explore this idea, we presented a simple step-stimulus with UV-light varying from 0% to 100% contrast around a mean background of 50% contrast (Fig. 5d, Vid. S4). On every other repetition, this UV-stimulus was superimposed on a naturalistic red-green-blue (RGB) background based on previous measurements of the spectrum of light in the zebrafish natural habitat (Methods). Finally, at the end of n=5 complete cycles, we presented a single, very bright UV-light flash to drive calcium to its light-evoked minimum. From here, we computed each UV-cone’s full dynamic range as the SyGCaMP6f-signal difference between the periods when all lights are off (maximal calcium) and when all lights are on (minimal calcium). Relative to this full dynamic range, we then computed each cone’s baseline during naturalistic stimulation when UV-light was held at 50% contrast. The resultant estimate of the calcium baseline across the eye recapitulated the previously observed brightness differences in the unstimulated eye: Signal baseline was maximal in the SZ, followed by a second, shallower peak around the nasal horizon (Fig. 5e, cf. Fig. 5c).

Next, we specifically compared response amplitudes to the 0% and 100% UV-contrast flashes during naturalistic background illumination in different zones. Like calcium baselines, this clearly showed that light and dark responses on average were most balanced in the SZ, followed by the nasal horizon, while both dorsal and ventral UV-cones were strongly dark biased (Fig. 5f).

To quantify this light-dark preference behaviour we calculated a Dark-Light-index (*DLi*) from each cone (Fig. 5g, see Methods), where a *DLi* of −1 indicates that a cone exclusively responds to the dark step, while a *DLi* of 1 corresponds to a fully light-biased response. A *DLi* of 0 denotes equal responsiveness to dark and light steps. This revealed that *DLi* varied with eye position, with the most balanced responses observed in the SZ and near the nasal horizon, recapitulating the previously observed gradual variations in calcium baseline (Fig. 5g, cf. Fig. 5c,e) and response properties (cf. Fig. 3f). In contrast, ventral and dorsal regions had a consistently negative DLi.

When compared directly, calcium baseline and DLi were strongly correlated (ρ = 0.85): a higher calcium baseline predicted a higher DLi (Fig. S2a-d). UV-cones from different eye-regions simply occupied different ranges of what appeared to be one continuum linking DLi and baseline. Taken together, our whole-eye imaging data therefore strongly suggests that systematic variations in calcium baseline are closely linked to a UV-cone’s preference for light or dark contrasts.

### Horizontal cells do not underlie functional differences in UV-cones

Differences in calcium baseline across UV-cones might be driven by differences in cone-intrinsic properties or by differential interactions with horizontal cells (HCs) (Chapot et al., 2017a; Van Hook et al., 2019; J. Klaassen et al., 2012; Thoreson and Mangel, 2012). In the latter case, variations in the strength of a tonic inhibitory input from HCs might drive variations in cone baseline and thus DLi. If this were the case, blockage of HC feedback should specifically elevate the low DLi of the dorsal and ventral retina. However, if anything, the opposite was observed. Pharmacological blockage of HCs using cyanquixaline (CNQX) did not elevate dorsal or ventral DLi, but instead slightly elevated DLi near the SZ and decreased it at the nasal horizon (Fig. S2e,f, Methods). Accordingly, it is unlikely that HCs strongly contribute to the observed functional differences in UV-cones. Instead, intrinsic differences in the properties of each UV-cone are likely dominant. What are these differences.

### Differential expression of phototransduction cascade genes is linked to calcium baseline changes

To pinpoint intrinsic differences between UV-cones that might underlie the observed elevation in calcium baseline leading to a light-preference in SZ UV-cones we used a transcriptomics approach (Stark et al., 2019). For this, we dissected entire retinas expressing GFP in all UV-cones and surgically separated the SZ from the remainder of the retina (non-SZ). We then dissociated and FACS-sorted UV-cones for subsequent transcriptomic profiling (Fig. 6a, Methods). Genes involved in phototransduction dominated the transcriptome of both SZ and non-SZ batches, with UV-opsin being the most strongly expressed protein-coding gene (Fig. 6b,c). Phototransduction genes were generally more highly expressed in SZ batches (Fig. 6d), consistent with their larger outer segment sizes (cf. Fig. 2). Accordingly, to compare the relative expression of key phototransduction genes, we normalized the expression level of each gene by the respective UV opsin expression level in each sample (Fig. 6e). This revealed that some key phototransduction genes had relatively higher expression in the SZ (e.g. gc3), while others were downregulated (e.g. cnga3 or gngt2b). Building on our exquisite understanding on phototransduction in general (Fain et al., 2010; Lamb, 2013; Pergner and Kozmik, 2017; Pugh and Lamb, 1993; Yau and Hardie, 2009), each of these regulatory changes can be quantitatively linked to a specific functional effect (Invergo et al., 2013, 2014). In particular, the tonic level of outer segment cGMP is key as it determines the open-probability of cyclic-nucleotide gated (CNG) channels which define the cone’s dark current to ultimately set the calcium baseline in the cone pedicle through the opening or closing of synaptic voltage gated calcium channels (Fain et al., 2010; Lamb, 2013; Pergner and Kozmik, 2017). Here, upregulation of guanylate cyclase 3 (Dizhoor and Hurley, 1999; Hurley, 1987; Kuhn, 2016) (gc3) is expected to boost the rate of cyclic guanosine monophosphate (cGMP) synthesis from guanosine triphosphate (GTP) to open CNG channels, depolarising the cone and thus rising synaptic calcium baseline. Similarly, downregulation of the gamma subunit of transducing (gngt2b) (Sprang et al., 2007) is expected to reduce the coupling between opsin excitation and phosphodiesterase 6 (pde6c) activation, thereby decreasing the rate of cGMP hydrolysis. This again will reduce CNG channel closing, depolarise the neuron and again elevate synaptic calcium.

**Figure 6.**
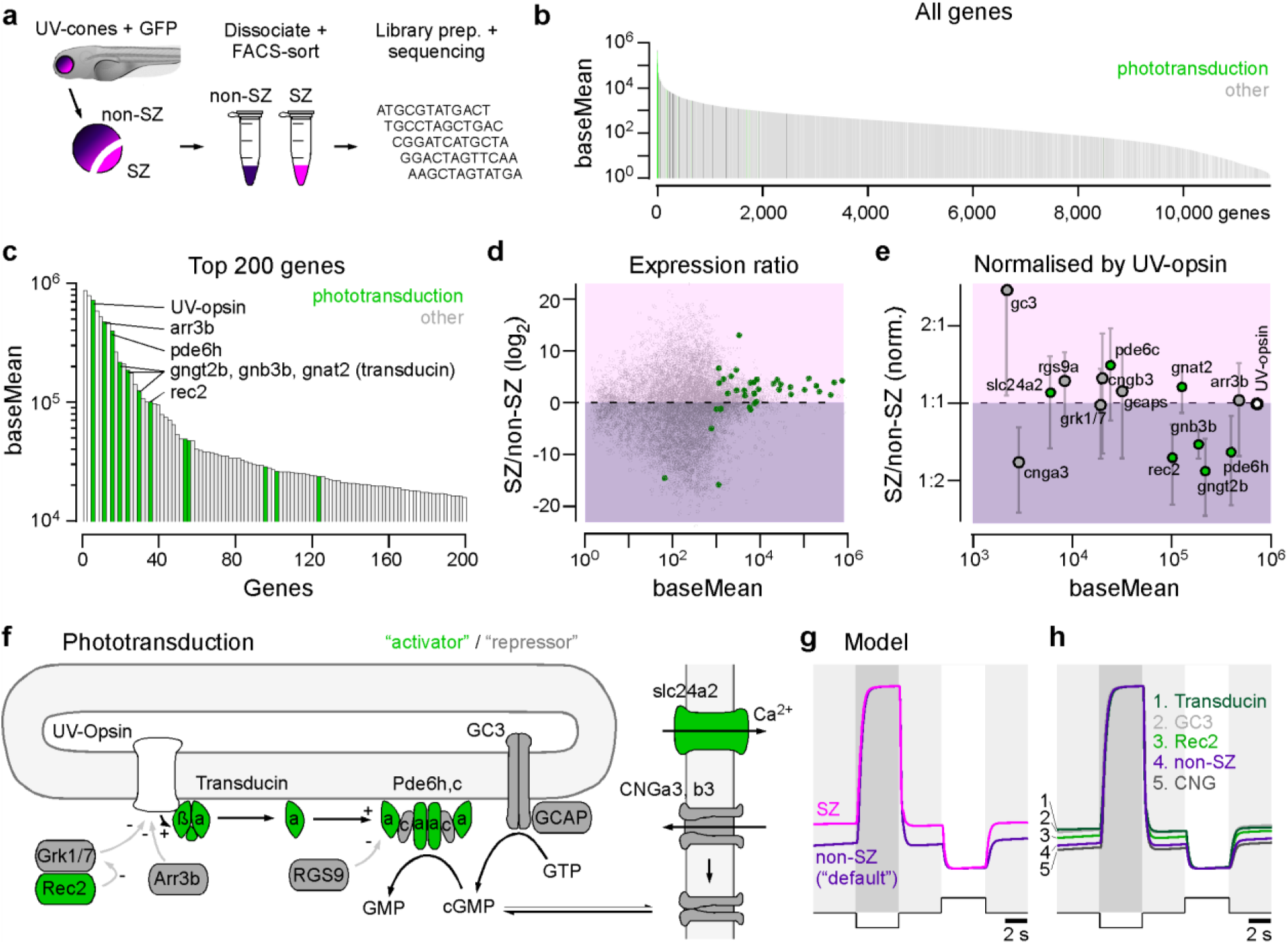
Tuning of phototransduction cascade elevates SZ baseline. **a**, UV-cone RNA-seq. workflow. Retinas from 7 dpf zebrafish Tg(opn1sw1:GFP) were dissected and separated into SZ and non-SZ. After cell dissociation, UV-cones were FACS-sorted and immediately flash frozen. Samples were then subjected to library preparation for next-generation sequencing. **b**, All detected genes in UV-cones ranked by expression label, with phototransduction genes highlighted and **c**, zoom in to the top 200 genes. The top 2 highly expressed genes are both non-protein coding genes, therefore UV-opsin is the highest expressed protein coding gene. **d**, Mean gene expression ratio between SZ and non-SZ batches, with phototransduction genes highlighted. **e**, as (d) but normalised to UV-opsin expression level in each batch and zoomed in to high expression phototransduction targets. Green and grey markers denote activators and repressors of the photo-response, respectively. Error bars in s.e.m.. **f**, Schematic of phototransduction based on ref (Yau and Hardie, 2009), with activators and repressors denoted in green and grey, respectively. **g**, Simulated current response of SZ and non-SZ UV-cones to 100% dark- and light-contrasts from a 50% contrast background based on ref (Invergo et al., 2014). Non-SZ was based on default model parameters, while SZ uses relatively scaled parameters according to gene expression ratios as in (e). **h**, Effects of expression changes of individual phototransduction components compared to non-SZ.

To quantitatively explore how these sum of all relative gene expression changes might affect the interplay of activators and repressors of the phototransduction cascade (Hurley, 1987; Pugh and Lamb, 1993; Pugh et al., 1999), we used a computational model of phototransduction in ciliary photoreceptors (Invergo et al., 2013, 2014) (Fig. 6f). We kept all pre-set parameters of the model constant, and only adjusted the relative levels of phototransduction elements according to the observed expression differences between SZ and non-SZ batches. In this way, we tested if we could turn a non-SZ cone (default model) into a SZ-cone through specific regulatory manipulations. Indeed, altering only the top four most differentially expressed targets (Transducin, GC3, Rec2 and CNG) phenomenologically reproduced the elevation in constitutive baseline and corresponding increase in the amplitude of the light-response in SZ cones (Fig. 6g). Already the modulation of single gene products’ relative expression levels could have a major effect, most notably in the case of transducin and GC3, and to a lesser extent also the downregulation of recoverin 2 (Rec2 (Dizhoor et al., 1991; Zang et al., 2015)) (Fig. 6h). Interestingly, transducin expression is also systematically adjusted between peripheral and foveal L/M-cones in the primate (Peng et al., 2019), indicating that this might constitute a regulatory hotspot for tuning cone function.

Nevertheless, not all activators of phototransduction were downregulated, and not all repressors were upregulated. For example, the downregulation of cnga3, a subunit of CNG channels, in the SZ had a minor hyperpolarising effect on cone-baseline. However, this was readily compensated for by the combination of the other expression differences. Indeed, the additional inclusion of all observed regulatory changes beyond the first four had a negligible effect on model baseline (not shown). Finally, to what extent additional changes might occur at targets with lower expression, such as synaptic calcium channels, remains an open question.

Taken together, our transcriptomics data therefore strongly suggests that a substantial fraction of the observed differences in light- over dark-preference in UV-cones across regions in the eye (cf. Figs. 3,5) have an origin in the regulation of phototransduction. In this view, the simultaneous up- and down-regulation of key phototransduction repressors and activators, respectively, will result in an increased dark current in SZ UV-cones and thereby boost constitutive opening of voltage activated calcium channels in the synapse, thus elevating the calcium baseline.

### Imaging synaptic release from cones *in vivo*

We next asked if and how the observed variations in UV-cone synaptic calcium are translated into rates of light-driven synaptic vesicle release in the live eye. Cones use ribbon synapses to release glutamate onto the dendrites of BCs and HCs (Baden et al., 2013a; Heidelberger et al., 2005; Lagnado and Schmitz, 2015; Moser et al., 2019; Regus-Leidig and Brandstätter, 2012; Sterling and Matthews, 2005; Thoreson, 2007) (Fig. 7a,b). Different ribbon designs are used extensively in sensory systems to support both transient and continuous high-throughput vesicle release with a broad spectrum of properties (Baden et al., 2013a; Bellono et al., 2018; Heidelberger et al., 2005; Lagnado and Schmitz, 2015; Wichmann and Moser, 2015). Accordingly, we wondered if UV-cones may also differentially tune their synaptic release machinery to further enhance the differences already present at the level of calcium and thus amplify eye-position specific signalling.

**Figure 7.**
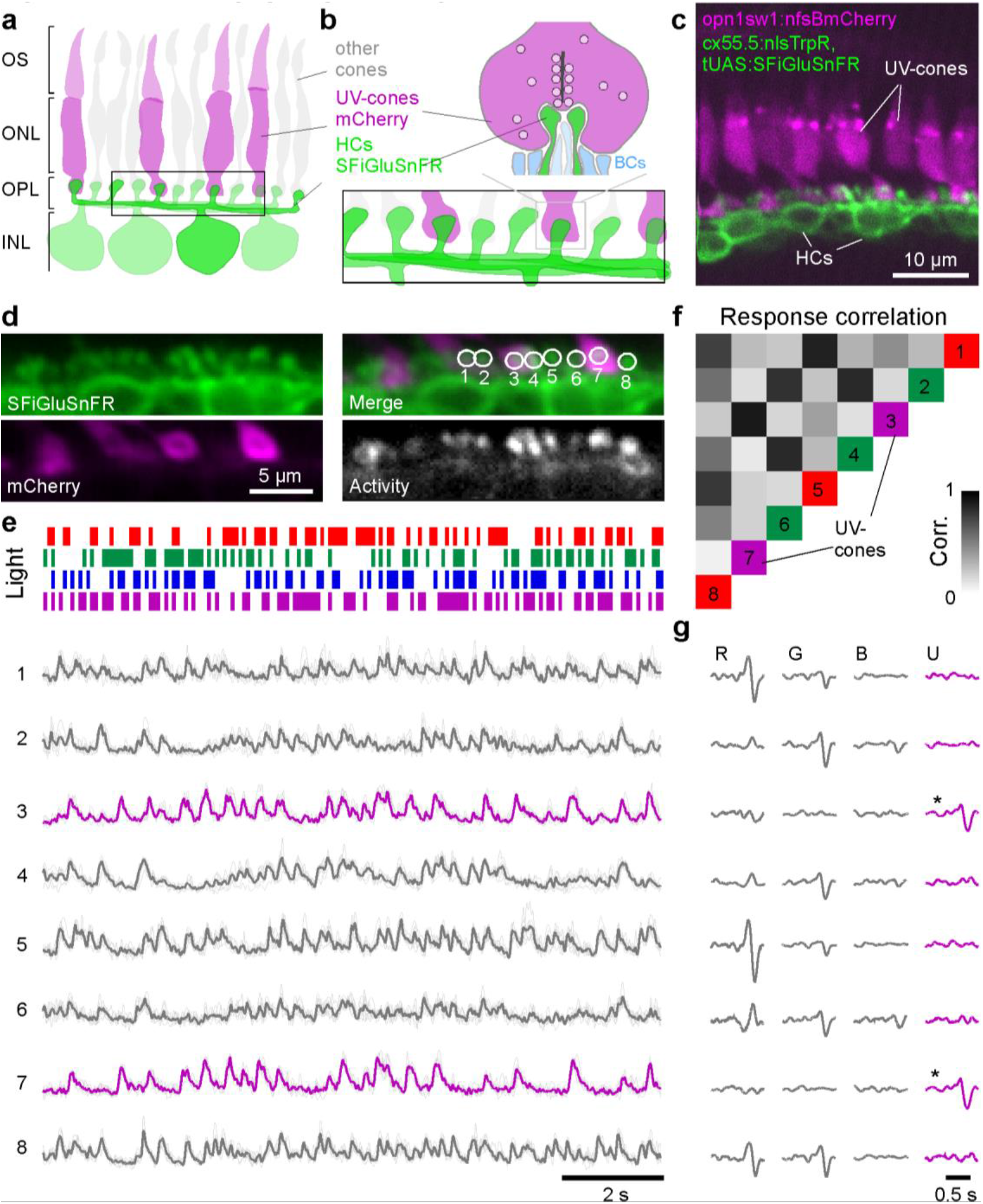
In vivo imaging of light-driven glutamate release from UV-cones. **a,b**, Schematic of HC dendrites at photoreceptor synaptic invaginations. SFiGluSnFR expression in HC dendrites is well-positioned to detect glutamate release from ribbon synapses (bar structure) at single terminals of any cone-type. UV-cones are identified by co-expression of mCherry as before. Outer segment (OS), outer nuclear layer (ONL), outer plexiform layer (OPL), inner nuclear layer (INL). **c**, In vivo two-photon image of SFiGluSnFR in HCs and nfsBmCherry in UV-cones. **d**, scan field for SFiGluSnFR recordings. Individual HC dendritic bundles at single cone terminals are readily visible. ROI 3 and 7 are associated with UV-cones as seen by overlap with the mCherry signal. A map of pixel-to-pixel correlation over time (Franke et al., 2017) highlights localised activity at each cone terminal. e, Exeat of mean and individual trial glutamate responses of ROIs from (d) to a tetrachromatic binary noise stimulus (Methods). UV-cone responses highlighted in magenta. **f**, Correlation of glutamate responses across pairs of ROIs. ROIs 3 and 7 are highly correlated only to each other. Colour code based on each ROIs preferred response as in (g). **g**, Linear filters (“kernels”) recovered by reverse correlation of each ROI’s response to the noise tetrachromatic stimulus (e): R, G, B, U denote red, green, blue and UV light, respectively. UV-cones are highlighted by *.

To address this question, we established optical glutamate recordings from single cones in the live eye by expressing the fluorescent glutamate biosensor SFiGluSnFR (Marvin et al., 2018) in postsynaptic HCs. HCs contact cones at specialised invaginations that are tightly sealed against the surrounding extracellular matrix (Chapot et al., 2017b; Regus-Leidig and Brandstätter, 2012), meaning that their dendrites can act as highly specific and spatially restricted glutamate antennas (Chapot et al., 2017b). As a population, HCs contact all four types of cones in the zebrafish eye (Klaassen et al., 2016; Li et al., 2009; Yoshimatsu et al., 2014), meaning that only a subset of HC dendritic signals correspond to synaptic release from UV-cones. To identify these contacts, we co-expressed mCherry in UV-cones (Fig. 7a-c). In an example recording from the nasal retina, we presented a 12.8 Hz tetrachromatic binary noise stimulus (Zimmermann et al., 2018) (Methods) and recorded the glutamate signals from the HC dendrites that innervate a row of neighbouring cones (Fig. 7d, Vid. S5). Amongst eight example regions-of-interest (ROIs), each covering a presumed single cone’s output site, two were identified as UV-cones based on mCherry co-expression (ROIs 3 and 7). Across glutamate responses within all eight ROIs (Fig. 7e), example sections of traces extracted for the two UV-cones were very similar to each other, but distinct from all other traces (Fig. 7e,f). Moreover, reverse correlation of each ROIs’ response to the noise stimulus revealed a pronounced UV-component for the two UV-cones, but diverse non-UV components in all other sites (Fig. 7g). This strongly indicated that there was no glutamate spill-over between neighbouring ROIs (see also discussions in refs (Chapot et al., 2017b; Franke et al., 2017; James et al., 2019)). Our approach therefore allowed recording UV-cone driven glutamate in the live eye at single pedicle resolution. We next used this approach to compare UV-cones’ calcium-to-glutamate transfer functions in different parts of the eye.

### Glutamate-release accentuates existing differences in presynaptic calcium

In nature, photoreceptors are constantly exposed to a rapidly changing stream of light- and dark-events as animals explore their visual environment. To explore how UV-cones in different parts of the eye differentially encode complex light-dark sequences, we recorded calcium and glutamate responses to the tetrachromatic binary noise-stimulus. Superimposition of the average calcium (top) and glutamate (bottom) responses to this stimulus from SZ and dorsal UV-cones revealed a marked difference in the synaptic transfer between these zones (Fig. 8a, Vid. S6): Despite relatively similar responses at the level of calcium (top), only SZ UV-cones responded strongly to the most rapid of stimulus reversals (bottom, arrowheads). These differences, which could not be explained by differences in the kinetics of GCaMP6f (Chen et al., 2013) and SFiGluSnFR (Marvin et al., 2018) (Fig. S3a), were subtly visible at the level of calcium, but they were strongly accentuated at the level of glutamate. Qualitatively similar effects were observed across all four zones (Fig. S3b-e).

**Figure 8.**
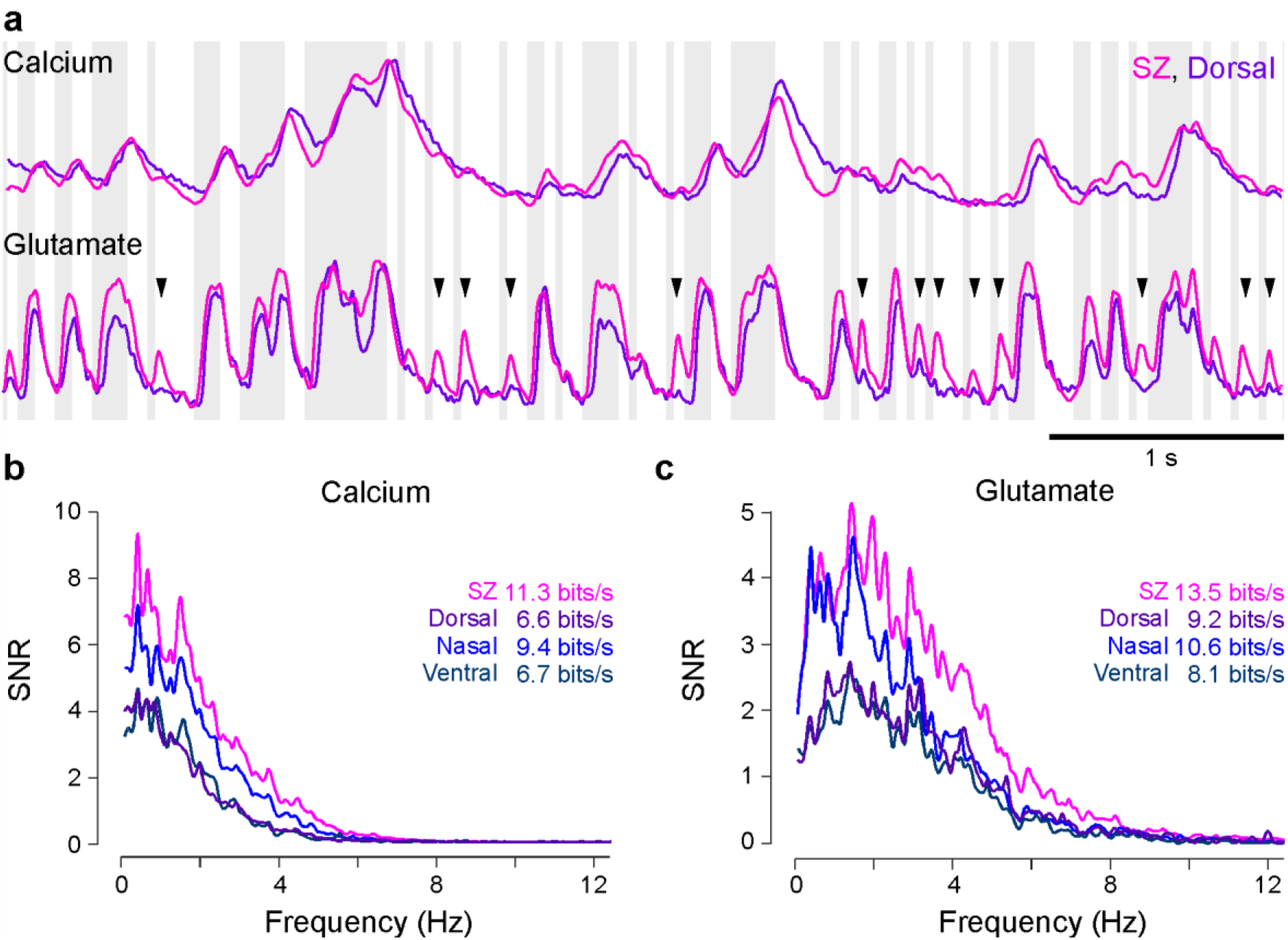
Synaptic release accentuates functional differences between UV-cones. **a,** Exeat of mean calcium (SyGCaMP6f) and glutamate (SFiGluSnFR) responses of SZ and dorsal UV-cones to the tetrachromatic noise stimulus (cf. Fig. 7e). Background shading indicates UV-light and dark stimulus periods. Arrowheads highlight enhanced glutamate response transients from SZ relative to dorsal UV-cones. **b**, Signal to noise ratio in the Fourier-domain and resulting information rate in calcium responses across UV-cones from different regions. **c**, as (b), computed for glutamate responses. n = 35, 20, 28, 18 for calcium in SZ, D, N and V, respectively and 51, 20, 22, 18 for glutamate SZ, D, N and V, respectively.

To determine how these differences can be linked to the amount of information which can be linearly decoded from the responses in each case, we computed the information rate based on the signal to noise ratio (SNR) in the Fourier domain (Fig. 8b,c)(van Hateren and van der Schaaf, 1998). Both at the level of calcium and glutamate, the most linearly decodable information was found in SZ-cones, followed by nasal, dorsal and finally ventral UV-cones. Moreover, the more pronounced glutamate responses of the SZ to rapid stimulus changes led to a higher SNR especially in the higher frequency domain. Accordingly, our glutamate imaging experiments clearly demonstrated that the differential tuning of UV-cones at the level of anatomy (Fig. 2), phototransduction (Fig. 6) and synaptic calcium (Figs. 3-5) is further enhanced at the level of synaptic release (Figs. 7, 8). Taken together, SZ UV-cones are therefore tuned to best support prey capture behaviour through the intricate adjustment of multiple mechanisms that involve all physiologically critical parts of the cell, from initial photon absorption in the outer segment to vesicle release at the synapse.

## DISCUSSION

We have shown that larval zebrafish must use single UV-cones at a time to detect the UV-bright microorganisms they feed on (Figs. 1, 4). For this, UV-cones in the retina’s strike zone are particularly dense and exhibit grossly enlarged outer segments (Fig. 2, Fig. S1) to boost local UV-photon detection efficiency. This is complemented by an elevation in these UV-cones’ synaptic calcium baseline (Figs. 3, 5) that likely stems from molecular retuning of the phototransduction machinery (Fig. 6) rather than interactions with horizontal cells. This leads to an increased dynamic range for encoding UV-bright events (Fig. 3) and sets of the capacity for increased information transfer across the synapse at the level of vesicle release driving retinal circuits (Figs 7,8). Together, UV-cones specifically in the strike zone are therefore exquisitely tuned to support the visual detection of prey. In contrast the remainder of the UV-detector array is less dense, uses smaller outer segments and a lower calcium baseline to detect large UV-dark objects, such as predators. In doing so, non-SZ UV-cones signal more sparsely and presumably conserve energy.

### Studying prey capture behaviour in the lab

Larval zebrafish prey capture behaviour has been extensively studied in the lab (Bianco et al., 2011; Gahtan, 2005; Jouary et al., 2016; McElligott and O’Malley, 2005; Muto and Kawakami, 2013; Novales Flamarique, 2016; Patterson et al., 2013; Semmelhack et al., 2014; Trivedi and Bollmann, 2013), though never specifically using UV-light. Nevertheless, even under these low-UV conditions, zebrafish do perform the behaviour. This suggests that non-UV cones feed into prey-capture circuits, perhaps to boost signal power in the absence of systematic background clutter, as is also the case under typical lab conditions. In support, the strong UV-dominance in SZ BCs is complemented small signals elicited also at other wavelengths, most notably in the blue range of the spectrum (Zimmermann et al., 2018). In parallel, it is important to consider the specific absorption spectrum of the zebrafish UV-opsin relative to the spectrum of any illuminating light. For safety reasons, commercially available TFT monitors (Franke et al., 2019), projectors and organic-LED (OLED) screens used in behavioural experiments tend to restrict short wavelengths to <1% signal power below 420 nm. In contrast, zebrafish UV-opsin absorption peaks at 365 nm (Chinen et al., 2003; Robinson et al., 1993), meaning that the short-wavelength signal of most of these light sources will activate the UV-opsin with <1% efficiency. Nevertheless, owing to the extreme photon-catch efficiency of SZ-UV cones this might still generate a small signal, provided the screen is sufficiently bright. Moreover, different projection set-ups (Bianco et al., 2011; Jouary et al., 2016; Semmelhack et al., 2014; Trivedi and Bollmann, 2013) or live paramecia illuminated by indoor lighting (Bianco et al., 2011; Gahtan, 2005; McElligott and O’Malley, 2005; Patterson et al., 2013) or indeed a fluorescence microscope’s excitation light (Muto et al., 2013) might afford higher spectral overlap. In the future it will therefore be critical to establish in more detail how the addition of UV-light affects behavioural performance.

### Mechanisms of photoreceptor tuning in vertebrates

For all we know, all sighted vertebrates have at least a mild form of an *area temporalis* or an *area centralis*, and in some species such as many primates as well as birds of prey and species of reptiles and fish, these specialised regions have further evolved into a fovea (Bringmann, 2019; Bringmann et al., 2018; Collin et al., 2000; Land, 2015). However, data on the possibility of regional tuning of photoreceptor function across most of these species remains outstanding with the notable exception of primates (Baudin et al., 2019; Sinha et al., 2017), mice (Baden et al., 2013b), and now zebrafish. In each of these latter three, cone-function has been found to be regionally tuned.

In many ways both the ‘purpose’ of functional tuning of SZ UV-cones as well as the underlying cellular and molecular mechanisms are reminiscent of differences between peripheral and foveal cones of the primate retina (Baudin et al., 2019; Curcio et al., 1990; Kemp et al., 1988; Mowat et al., 2008; Peng et al., 2019; Sinha et al., 2017). For example, in both zebrafish SZ UV-cones and in primate foveal cones outer segments are elongated (Curcio et al., 1990; Packer et al., 1989) and light-response kinetics are slowed (Baudin et al., 2019; Sinha et al., 2017). In the primate fovea, expression of rod-transducin gamma subunit has been discussed as one determinant of the slowed kinetics (Peng et al., 2019), which conceptually links with our finding of reduced levels of cone-transducin gamma subunit in zebrafish SZ cones. In each case, these structural and functional alterations can be linked to an increased capacity for the detection of low numbers of photons and subsequent signal processing. In the primate fovea, they are critical to keep noise at bay to supply a low-convergence postsynaptic retinal network (Ala-Laurila et al., 2011; Angueyra and Rieke, 2013). Establishing to what extent the postsynaptic networks in the zebrafish’s SZ resemble those of the primate fovea will be an important area of research in the future. Nevertheless, already now it seems clear that noise-reduction will be an asset also for SZ UV-cones. In contrast to primates and zebrafish, mice have only a very mild area centralis aligned with visual space above the nose (Bleckert et al., 2014; Dräger and Olsen, 1981; Salinas-Navarro et al., 2009). However, they feature a pronounced opsin expression gradient across the retina’s dorsal-ventral axis (Szél et al., 2000) which has been linked to differential processing of light- and dark-contrasts (Baden et al., 2013b), much in line with observed differences in zebrafish UV-cones. However, unlike in zebrafish, ventral short-wavelength vision in mice is dark-biased (Baden et al., 2013b), which rather hints at the flexibility in how photoreceptors can be tuned to support specific visual tasks.

For the most part, the detailed cellular and molecular mechanisms that lead to differential cone-tuning across the retinal surface in mice and primates remain to be established. Building on our work, we anticipate that the possibility to perform high-throughput *in-vivo* experiments in genetically modified larval zebrafish will be a major asset for studying mechanisms of photoreceptor tuning in general.

### Synaptic tuning through the ribbon

Beyond altering the morphological and biochemical properties of the outer segment, our results further suggest that also the pedicle is functionally adjusted to support distinct modes of calcium-dependent vesicle release in UV cones in different parts of the eye. Cones use ribbon-type synapses which have been a key focus for investigating the functional tuning of neural circuits (Baden et al., 2013a; Bellono et al., 2018; Heidelberger et al., 2005; Lagnado and Schmitz, 2015; Moser et al., 2019; Regus-Leidig and Brandstätter, 2012; Sterling and Matthews, 2005; Thoreson, 2007; Wichmann and Moser, 2015). For example, electro-sensory ribbon synapses in rays and sharks are differentially tuned at the level of both synaptic ion channels and ribbon morphology to support the encoding distinct signal frequency bands required by these two groups of animals (Bellono et al., 2018). Indeed, ribbon synapses across species and modalities support a vast range of functional properties, and generally the structure and function of each group of synapses can be closely linked to specific signalling requirements (Heidelberger et al., 2005; Lagnado and Schmitz, 2015; Moser et al., 2019; Sterling and Matthews, 2005; Thoreson, 2007). While therefore ribbon synapses do strongly vary across distinct sets of neurons that support diverse functional tasks, to our knowledge this type of tuning has not been studied across a single neuron type. Accordingly, in the future it will be important to establish if and how the observed differences in synaptic transfer functions across zebrafish UV-cones can be linked to structural and molecular differences in the synapse itself.

### Retinal and central wiring for prey capture

The region-specific differences in UV-cone function present the first pre-processing steps to detect prey and predators already at the visual system’s first synapse (Baden et al., 2013b; Chapot et al., 2017b). However, how these signals are used by retinal and brain networks for robust extraction of such behaviourally crucial information remains an open question. Ultimately, light patterns picked up by distinct regions of the UV detector array must lead to the differential activation of brain circuits that control distinct behavioural programmes (Dunn et al., 2016; Preuss et al., 2014; Semmelhack et al., 2014). For this, the signal must first travel to the feature extracting circuits of the inner retina (Baden et al., 2019; Masland, 2001) via the diverse set of retinal bipolar cells (Connaughton and Maguire, 1998; Connaughton and Nelson, 2000; Li et al., 2012; Zimmermann et al., 2018). Previous work highlighted a strong dominance of inner retinal UV-On circuits specifically in the strike zone (Zimmermann et al., 2018), suggesting that the signal from SZ UV-cones is indeed selectively picked up by a subset of local UV-On BCs for further processing. Next, the UV-signal must be selectively sent to the specific relevant processing centres of the brain (Connaughton and Nelson, 2015; Robles et al., 2014; Xiao and Baier, 2007). In agreement, pretectal arborisation field 7 (AF7) which underpins prey-capture behaviour is mainly innervated by temporal but not nasal RGCs (Semmelhack et al., 2014), strongly hinting that AF7 may be predominately driven by SZ-circuits. Clearly, circuits for prey capture in larval zebrafish are both anatomically (Semmelhack et al., 2014) and functionally (Zimmermann et al., 2018) regionalised to drive a regionally biased behavioural repertoire (Bianco et al., 2011; Mearns et al., 2019). To what extent this can be supported through regional tuning of neuron-types alone – as in case of UV-cones - or in addition requires the specific positioning of unique neuron types in different parts of the eye and brain will be important to address in the future. Indeed, transcriptomic analysis recently highlighted the putative presence of one “extra” bipolar cell type specifically in the primate fovea (Peng et al., 2019), with yet unknown morphology and function.

## MATERIALS AND METHODS

### Imaging the appearance of paramecia at different wavelengths of light

*Paramecium caudatum* (Sciento, P320) were placed in a container filled with fish water and pebbles, to approximately mimic a zebrafish natural habitat (Zimmermann et al., 2018). Images were taken outdoors under the sun (typical sunny day in UK, Brighton in May, no cloud at around 1 pm) with a CCD camera (Thorlabs DCU223M) fitted with a lens (Thorlabs ACL1815L), a constitutive glass filter (Thorlabs FGB37) as well as switchable glass filters (UV: FGUV11-UV, Yellow: FGV9; both Thorlabs) on a filter-wheel. Videos were acquired at 10 Hz, with single frame exposure times of 1 and 70 ms for yellow and UV, respectively. The focal distance of the camera was ~2.5 cm, and it was positioned against the wall of the tank from the outside. The effective recording spectra were computed by multiplying the spectral sensitivity of the camera chip itself with all optical components in the path.

### Animals

All procedures were performed in accordance with the UK Animals (Scientific Procedures) act 1986 and approved by the animal welfare committee of the University of Sussex. For all experiments, we used *6-8 days post fertilization* (*dpf*) zebrafish (Danio rerio) larvae. The following previously published transgenic lines were used: *Tg(opn1sw1:nfsBmCherry)* (Yoshimatsu et al., 2016), *Tg(opn1sw1:GFP)* (Takechi et al., 2003). In addition, *Tg(opn1sw1:GFP:SyGCaMP6f)*, *Tg(cx55.5:nlsTrpR)*, and *Tg(tUAS:SFiGluSnFR)* lines were generated by injecting pBH-opn1sw1-SyGCaMP6f-pA, pBH-cx55.5-nlsTrpR-pA (Yoshimatsu et al., 2016), or pBH-tUAS-SFiGluSnFR-pA plasmids into single-cell stage eggs. Injected fish were out-crossed with wild-type fish to screen for founders. Positive progenies were raised to establish transgenic lines.

All plasmids were made using the Gateway system (ThermoFisher, 12538120) with combinations of entry and destination plasmids as follows: pTo2pA-opn1sw1-SyGCaMP6f-pA: pTol2pA (Kwan et al., 2007), p5E-opn1sw1 (Yoshimatsu et al., 2016), pME-SyGCaMP6f, p3E-pA (Kwan et al., 2007); pBH-tUAS-SFiGluSnFR-pA: pBH (Yoshimatsu et al., 2016), p5E-tUAS (Suli et al., 2014), pME-SFiGluSnFR, p3E-pA. Plasmid pME-SyGCaMP6f was generated by inserting a polymerase chain reaction (PCR)-amplified GCaMP6f (Chen et al., 2013) into pME plasmid and subsequently inserting a PCR amplified zebrafish synaptophysin without stop codon at the 5’ end of GCaMP6f. pME-SFiGluSnFR was made by inserting a PCR amplified SFiGluSnFR (Marvin et al., 2018) fragment in pME plasmid.

Animals were housed under a standard 14:10 day/night rhythm and fed three times a day. Animals were grown in 0.1 mM 1-phenyl-2-thiourea (Sigma, P7629) from 1 *dpf* to prevent melanogenesis. For 2-photon *in-vivo* imaging, zebrafish larvae were immobilised in 2% low melting point agarose (Fisher Scientific, BP1360-100), placed on a glass coverslip and submerged in fish water. Eye movements were prevented by injection of a-bungarotoxin (1 nL of 2 mg/ml; Tocris, Cat: 2133) into the ocular muscles behind the eye. For some experiments, CNQX (~0.5 pl, 2 mM, Tocris, Cat: 1045) in artificial cerebro-spinal fluid (aCSF) was injected into the eye.

### Behavioural experiments

Individual 7-8 *dpf* zebrafish larva were head-mounted in 2% low-melting-point agarose (Fisher Scientific, BP1360-100) in a 35 mm petri dish with the eyes and tail free to move and filmed under infrared illumination (940 nm) using a Raspberry Pi camera at 30 Hz based on a previous design (Maia Chagas et al., 2017). An Arduino-microcontroller was used to iteratively switch top-illumination of the dish between UV (374±15 nm) or red (620±10 nm) LED light in periods of 2 minutes. The same fish was filmed continuously for three such cycles (total of 12 minutes per n=12 fish), and behavioural performance was manually annotated offline as either a “full prey capture bout” (eye convergence plus tail movement) or “tracking” (single or bilateral eye movements in the absence of tail movements).

### UV-cone density estimation across the visual field

The UV-cone distribution across the eye was first established from confocal image stacks of Tg(opn1sw1:GFP) eyes from *7 dpf* larvae where all UV-cones are labelled. Fish were mounted with one eye facing the objective lens. As in previous work (Zimmermann et al., 2018) the locations of all UV-cones in the 3D eye were detected using a custom script in Igor Pro 6.3 (Wavemetrics). To project the resultant UV-cone distribution into visual space, we first measured the eye size as being 300 μm on average. In addition, we determined that both the eyeball and the lens follow a nearly perfect spherical curvature with a common point of origin. From this, we assumed that any given UV-cone collects light from a point in the space that aligns with a straight line connecting the UV-cone to the outside world through the centre of the lens. From here, we mapped UV-cone receptive field locations across the full monocular visual field.

### Immunostaining, dye-staining and confocal imaging

Larval zebrafish (*7-8 dpf*) were euthanised by tricane overdose and then fixed in 4% paraformaldehyde (PFA, Agar Scientific, AGR1026) in PBS for 30 min at room temperature. After three washes in PBS, whole eyes were enucleated and the cornea was removed by hand using the tip of a 30 G needle. Dissected and fixed samples were treated with PBS containing 0.5% TritonX-100 (Sigma, X100) for at least 10 mins and up to 1 day, followed by the addition of primary antibodies. After 3-5 days incubation at 4°C, samples were washed three times with PBS 0.5% TritonX-100 solution and treated with secondary antibodies and/or BODIPY (ThermoFisher, D3835) dye. After one day incubation, samples were mounted in 1% agar in PBS on a cover slip and subsequently PBS was replaced with mounting media (VectaShield, H-1000) for imaging. Primary antibodies used were anti-GFP (abcom, chicken, ab13970) and anti-CoxIV (abcom, rabbit, ab209727). Secondary antibodies were Donkey CF488A dye anti-chick (Sigma, SAB4600031) and Goat Alexa647 dye anti-rabbit (ThermoFisher, A-21244). Confocal image stacks were taken on a TSC SP8 (Leica) with 40x water immersion objective (C PL APO CS2, Leica), a 63x oil immersion objective (HC PL APO CS2, Leica) or a 20x dry objective (HC PL APO Dry CS2, Leica). Typical voxel size was 150 nm and 1 μm in xy and z, respectively. Contrast, brightness and pseudo-colour were adjusted for display in Fiji (NIH). Quantification of outer segment lengths and anti-GFP staining intensity was performed using custom scripts in Igor Pro 6.3 (Wavemetrics) after manually marking outer segment outer and inner locations.

### 2-photon calcium and glutamate imaging and light stimulation

All 2-photon imaging was performed on a MOM-type 2-photon microscope (designed by W. Denk, MPI, Martinsried; purchased through Sutter Instruments/Science Products) equipped with a mode-locked Ti:Sapphire laser (Chameleon Vision-S, Coherent) tuned to 927 or 960 nm for SyGCaMP6f and SFiGluSnFR imaging and 960 nm for mCherry and SFiGluSnFR double imaging. We used two fluorescence detection channels for SyGCaMP6f/iGluRSnFR (F48×573, AHF/Chroma) and mCherry (F39×628, AHF/Chroma), and a water immersion objective (W Plan-Apochromat 20x/1,0 DIC M27, Zeiss). For image acquisition, we used custom-written software (ScanM, by M. Mueller, MPI, Martinsried and T. Euler, CIN, Tuebingen) running under IGOR pro 6.3 for Windows (Wavemetrics).

Recording configurations were as follows: SyGCaMP6f UV flashes Fig. 3: 128×16 pixels (1 ms per line, 62.5 Hz); SyGCaMP6f whole-eye Fig. 5: 512×512 pixels (2 ms per line, 0.97 Hz), SFiGluSnFR noise recording Fig. 7: 128×32 pixels (1 ms per line, 31.25 Hz), SFiGluSnFR and SyGCaMP6f noise recordings Fig. 8: 64×4 pixels (2 ms per line, 125 Hz). Light stimulation was setup-up as described previously (Zimmermann et al., 2018). In brief, light stimuli were delivered through the objective, by band-pass filtered light emitting diodes (LEDs) (‘red’ 588 nm, B5B-434-TY, 13.5cd, 20 mA; ‘green’ 477 nm, RLS-5B475-S; 3-4 cd, 20mA; ‘blue’ 415 nm, VL415-5-15; 10-16 mW, 20 mA; ‘ultraviolet, UV’ 365 nm, LED365-06Z; 5.5 mW, 20 mA, Roithner, Germany). LEDs were filtered and combined using FF01-370/36, T450/pxr, ET420/40 m, T400LP, ET480/40x, H560LPXR (AHF/Chroma) and synchronized with the scan retrace at 500 (2 ms lines) or 1,000 Hz (1 ms lines) using a microcontroller and custom scripts (available at https://github.com/BadenLab/Zebrafish-visual-space-model). The ratio of LED intensities was calibrated (in photons per s per cone) such that each LED would relatively stimulate its respective cone-type as it would be activated under natural spectrum light in the zebrafish habitat (Zimmermann et al., 2018): 34, 18, 4.7 and 2.1 x10^5^ photons per cone per s for red-, green-, blue-, and UV-cones, respectively. We used these “natural spectrum” LED intensities as a background light and modulated contrasts depends on experiments. LED contrasts were 0% for dark and 2,500% for bright flashes (Fig. 3b-f), 0% background and 2,500% flash (Fig. 3g,h), 2,500% background and 0% dark flash (Fig. 3i,j), 0% dark and 200% bright (Fig. 5). For tetrachromatic noise (Figs. 7-8), each of 4 LEDs was simultaneously but independently presented at 100% contrast in a known sequence at 12.8 Hz. For all experiments, the animal was kept at constant background illumination for at least 5 s at the beginning of each recording to allow for adaptation to the laser.

### UV-cone activation model

Cone distributions were taken from published data (Zimmermann et al., 2018). UV- and blue-cones were taken from the same representative eye and aligned with red- and green-cones from a second eye and projected into visual space. The full array was cropped at ±60°. Model BCs were randomly spaced at a minimum radius of 10°. BCs summed the activity from all cones within this same fixed radius. Target trajectory was computed as a random walk on an infinite plane (canonical diffeomorphism), such as the left/right and top/bottom borders are continuous with each other. At each 1° step-size iteration (equivalent to 10 ms), the target advanced at a constant speed of 100°/s with a random change of angle (α) that satisfied - 15° < α < 15°. Cone activation by the moving target was computed as follows: At each time-point, the distance between the centres of the target and each cone was determined. If this distance was smaller than the sum of the target radius (1° and 2.5° for light and dark target, respectively) and a cone’s receptive field radius (0.38°), the cone was activated to yield a binary activation sequence over time for each cone. This sequence was then convolved with the cone’s impulse response. Here, the peak amplitude and recovery time constant was assigned based on a cone’s position, drawing on the four measurement points established from calcium imaging (dorsal, nasal i.e. horizon, ventral and SZ, cf. Fig 3). Along the dorsal-ventral axis, values were chosen based on the relative distance between the horizon and the dorsal or ventral edge. For example, a cone positioned 75% towards the dorsal edge from the horizon would be assigned values weighted as 0.75:0.25:0 from dorsal, nasal and ventral measurements, respectively. In addition, if a cone was within 30° of the SZ centre (−30°,−30°), it was in addition weighted based on values from the SZ in the same way. In each run, all activation values were normalised to the peak activation across the entire array.

### RNA-sequencing of UV-cones

Whole *7 dpf Tg(opn1sw1:GFP*) larval zebrafish retinas were dissected in carboxygenated aCSF (CaCl 0.1275 g/L, MgSO_4_ 0.1488 g/L, KCl 0.231 g/L, KH_2_PO_4_ 0.068 g/L, NaCl 7.01 g/L, D-Glucose 1.081 g/L, and NaHCO_3_ 1.9 g/L) while keeping track of each retina’s orientation. Each retina was then cut into two pieces: SZ, and non-SZ. Typically tissues from ~10 fish (20 eyes) were batched into one tube and dissociated using a papain dissociation system (Worthingtonm LK003176, LK003170, LK003182) with the following modification in the protocol: Incubation in papain for 10 min at room temperature. During dissociation, tissues were gently pipetted every 3 min to facilitate dissociation using glass pipette with rounded tip. After 10 min incubation, DNase and ovomucoid were added and the tissues were further mechanically dissociated by gentle pipetting. Dissociated cells were immediately sorted for GFP expression by FACSMelody (BD Biosciences). Approximately 100 cells were sorted in one tube, flash frozen in liquid nitrogen and stored at −80 degree until further use. Libraries were prepared using Ultra-low input RNA kit (Takara, 634888) and subjected to next generation sequencing at GENEWIZ (NZ, US). Sequencing data was quality checked and trimmed to remove adaptors using Trim Galore!([CSL STYLE ERROR: reference with no printed form.]), aligned on the zebrafish genome (GRCz11.9) in HISAT2 (Kim et al., 2015), and counted for gene expression in featureCounts (Liao et al., 2014) using the public server at the usergalaxy.org online platform (Afgan et al., 2018). In total, four repeats each were performed for SZ and non-SZ samples.

### Differential gene expression analysis

For the analysis of differential gene expression of the SZ vs non-SZ we used the DESeq2 package in R/Bioconductor (Love et al., 2014). We only included genes which had a count of at least 5 sequence fragments in at least 2 of the 8 samples (4 SZ + 4 non-SZ). Since we wanted to measure the effect between zones, controlling for differences in the individual eyes, we included the eye as an additional latent variable (design = ~eye+zone). The DESeq2 package then uses a generalised linear model with a logarithmic link to infer a negative-binomial distribution for gene counts (Love et al., 2014). The inferred means via the *poscount* estimator, which calculates a modified geometric mean by taking the n^th^ root of the product of non-zero counts, are shown in Fig. 6b,c. The log-fold changes (Fig.6d) were then also estimated in DESeq2.

For determining differential expression normalised by UV-opsin (Fig. 6e) we instead calculated using the raw count data, normalised by the count of the UV-opsin gene. From here, mean fold changes were calculated by taking fold changes of individual SZ and non-SZ sample pairs.

### Modelling phototransduction

We used a previously described and verified computational model of phototransduction in vertebrate ciliary photorecptors (Invergo et al., 2013, 2014). We simulated the photo-response to 100% dark or 100% bright contrasts using default parameters provided by the model for non-SZ simulation. For simulating the SZ, we then scaled all according to the relative gene expression change between SZ and nSZ conditions. Transducin was scaled by taking the lowest value among components (gngt2b, gnb3b, gnat2) because all components are necessary for transducin function. Similarly, we scaled CNG based on the CNGa3 expression level. Parameters changed for each condition are listed in Supplementary Table S1.

### Software

Data analysis was performed using IGOR Pro 6.3 (Wavemetrics), Fiji (NIH), Python 3.5 (Anaconda distribution, scikit-learn 0.18.1, scipy 0.19.0 and pandas 0.20.1) and R 3.5.1.

### Pre-processing and Dark-Light-index

Regions of interest (ROIs), corresponding to individual presynaptic terminals of UV-cones were defined automatically based on local thresholding of the recording stack’s s.d. projection over time (s.d. typically >25), followed by filtering for size and shape using custom written software on IGOR Pro 6.3 (Wavemetrics). Specifically, only round ROIs (<150% elongation) of size 2-5 μm^2^ were further analysed. For glutamate recording, ROIs were manually placed as the shape of HC dendritic terminals at cone terminals are often skewed. Calcium or glutamate traces for each ROI were extracted and z-normalised based on the time interval 1-6 s at the beginning of recordings prior to presentation of systematic light stimulation. A stimulus time marker embedded in the recording data served to align the traces relative to the visual stimulus with a temporal precision of 1 or 2 ms (depending on line-scan speed). The Dark-Light-index (DLi) was calculated as:

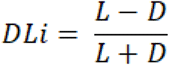

where *L* and *D* are the mode of response amplitudes to UV- and dark-flash with RGB background, respectively.

### Information Rates

To calculate information rates, we first filtered recorded traces for quality: We calculated the linear response kernel to UV-light stimulation for each trace and took only the traces where the response amplitude of the kernel, measured as its standard deviation, was at least 70% of the kernel with maximal response amplitude of the same zone.

We then followed the procedure as described in ref (Van Hateren and Snippe, 2001) using the bias correction method for finite data. For this, we assumed that the noise between repetitions of the experiment was statistically independent. For independent Gaussian statistics, the information rate *R* can be computed as:

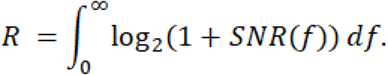

Since photoreceptors are best driven by low frequency signals (Baden et al., 2013a) we chose a cut-off frequency of 12 Hz. We then calculated a bias corrected signal to noise ratio (SNR) as:

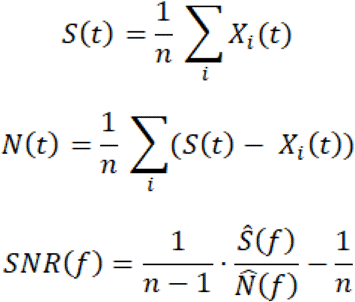

where *X_i_*, is an individual trial, *n* is the number of trials and 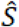 and 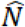 are the Fourier transform of *S* and *N*, respectively.

We used Welch’s method to reduce noise in the estimated power spectra.

### Statistics

No statistical methods were used to predetermine sample size. P-values were calculated using non-parametric Mann-Whitney U tests in Fig. 3h,j and Fig. S1g, and paired t-test in Fig. 1g. Owing to the exploratory nature of our study, we did not use randomization or blinding.

We used Generalized Additive Models (GAMs) to analyse the relationships between eye position and outer segment size, baseline, and dark-light index (Fig. 2g, Fig. 5c,e,g, Fig. S2e). GAMs can be seen as an extension to the generalized linear model by allowing linear predictors, which depend on smooth functions of the underlying variables(Wood). We used the mgcv-package (version 1.8-28) on a Windows 10 workstation (8 Xeon E3-1270 v5 3.6 GHz; 64 GB RAM) with default parameters. We modelled the dependence of the variable of interest as a smooth term with 20 degrees of freedom. In addition, we incorporated the fish id as a random effect. The models explained ~40-80% of the deviance. For plotting, we generated the predicted mean response with approximate 95% confidence intervals excluding fish id (this leads to a slight perceived offset between the raw data points and the mean response). Statistical significance for differences between the dependence of DLi in baseline and HC block conditions were obtained using the plot_diff function of the itsadug-package for R (version 2.3).

## Supporting information

SVideo 1

SVideo 2

SVideo 3

SVideo 4

SVideo 5

SVideo 6

STable 1

## Data and software availability

Preprocessed 2-photon imaging data, cone-density counts, natural imaging data and transcriptome data will be made freely available at http://www.badenlab.org/resources and http://www.retinal-functomics.net. All other data and code are available upon reasonable request.

## Author contributions

TY and TB designed the study, with input from CS and PB; TY generated novel lines and performed all data collection and pre-processing except behavioural experiments with input from TB; NEN performed and analysed behavioural experiments; TY and CS analysed transcriptomic data; CS computed information rates and deconvolutions. TY and PB performed statistical analysis. TY and TB wrote the manuscript with inputs from all authors.

## Acknowledgements

We thank Leon Lagnado, Thomas Euler and Julie Semmelhack for critical feedback. The authors would also like to acknowledge support from the FENS-Kavli Network of Excellence and the EMBO YIP.

