## Supplementary Data for "Cellular and molecular mechanisms of photoreceptor tuning for prey capture in larval zebrafish"

**SUPPLEMENTARY MATERIALS**

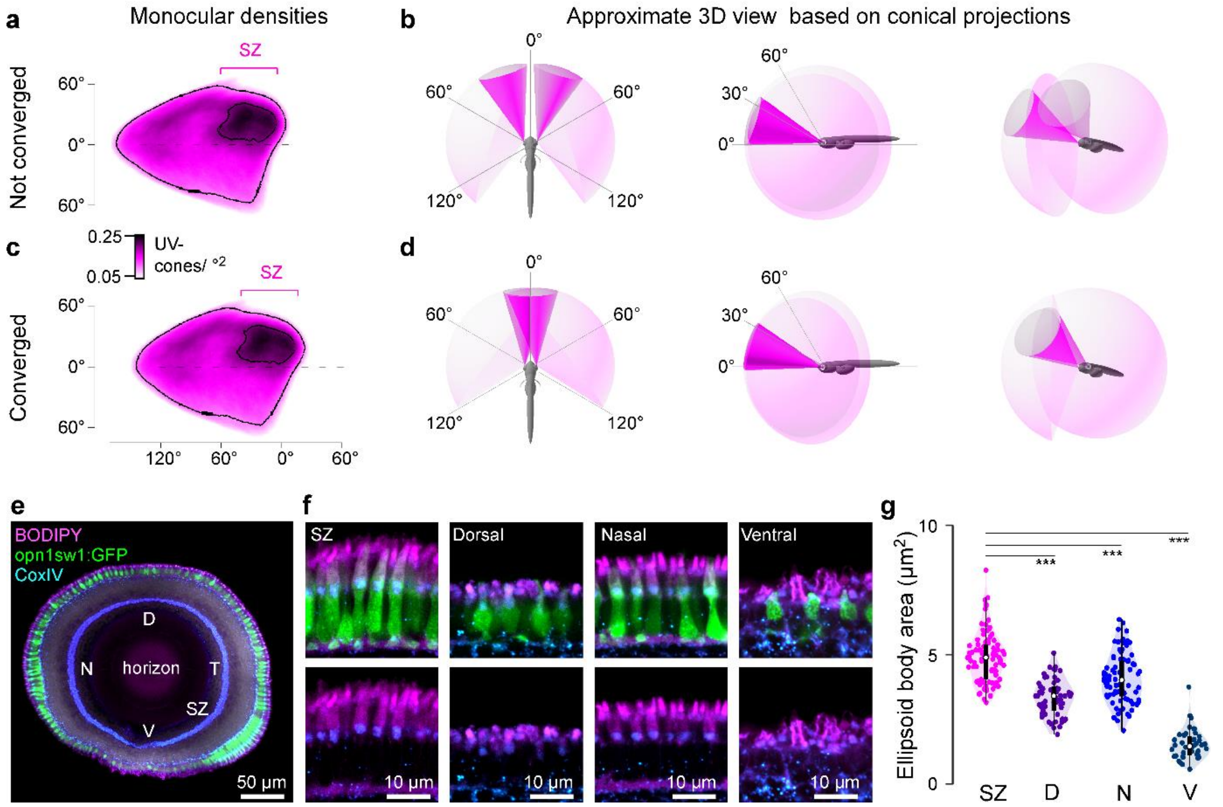


**Figure S1, related to Figure 2. Structural specialisations of UV-cones for prey capture.** **a**, Monocular UV-cone density projection into visual space when eyes are not converged. **b**, Schematics of approximate visual space surveyed by the two SZs (dark pink) and full field of view (light pink) when viewed from top (left), side (middle) and front/bottom (right). **c,d**, As (a,b), but when the eyes are converged. **e.** UV-cones (*Tg(opn1sw1:GFP)*) with BODIPY and mitochondria (CoxIV) counterstaining in a whole eye sagittal view. N, nasal; D, dorsal; T, temporal; SZ, strike zone; V, ventral. **f.** High magnification images of the same eye. **g**, Quantification of differences in ellipsoid body area between zones. Mann-Whitney U-test, ***: p<0.0001.

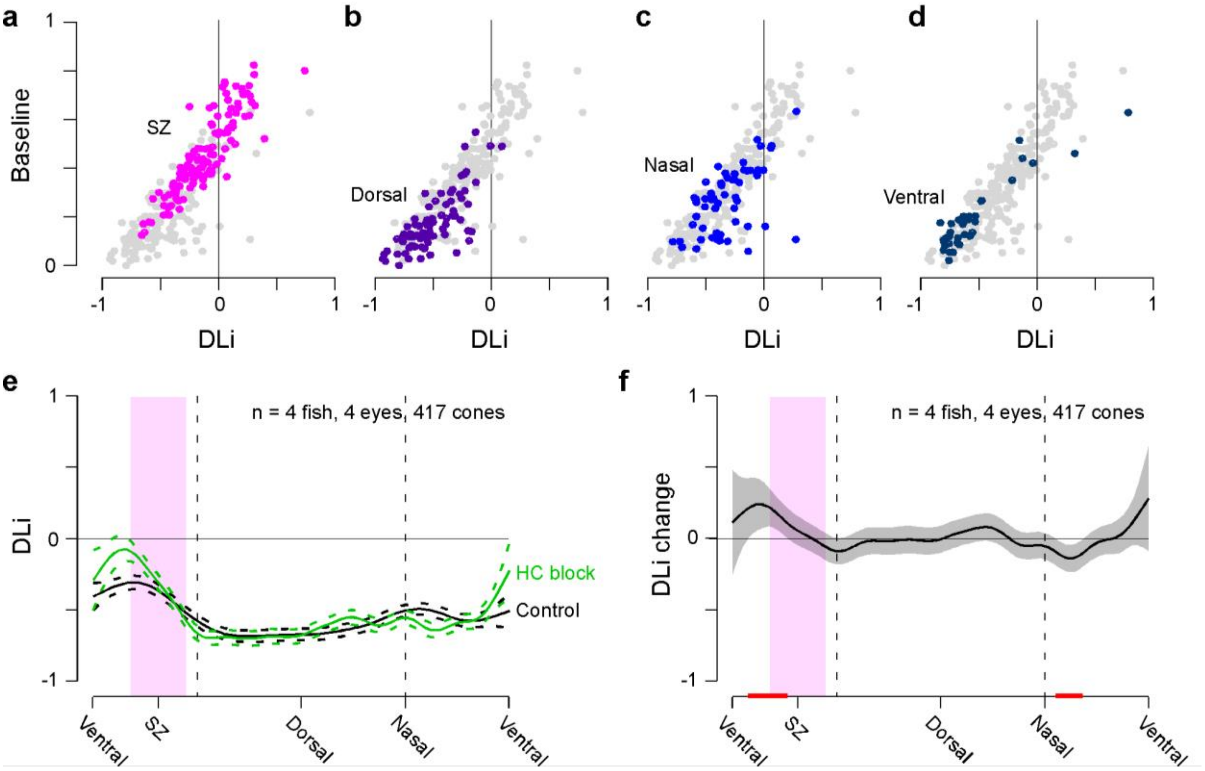


**Figure S2, related to Figure 5. Baseline relation to DLi and horizontal cell block. a-d**, Scatter plots of calcium baseline versus dark-light index (DLi) across zones, with full dataset (grey) superimposed by the individual zones as indicated. **e**, Mean and 95% confidence intervals of DLi before (black) and after (green) blockage of horizontal cell feedback by CNQX application. **f**, Change in DLi from (e), with red lines indicating significant change from 0.

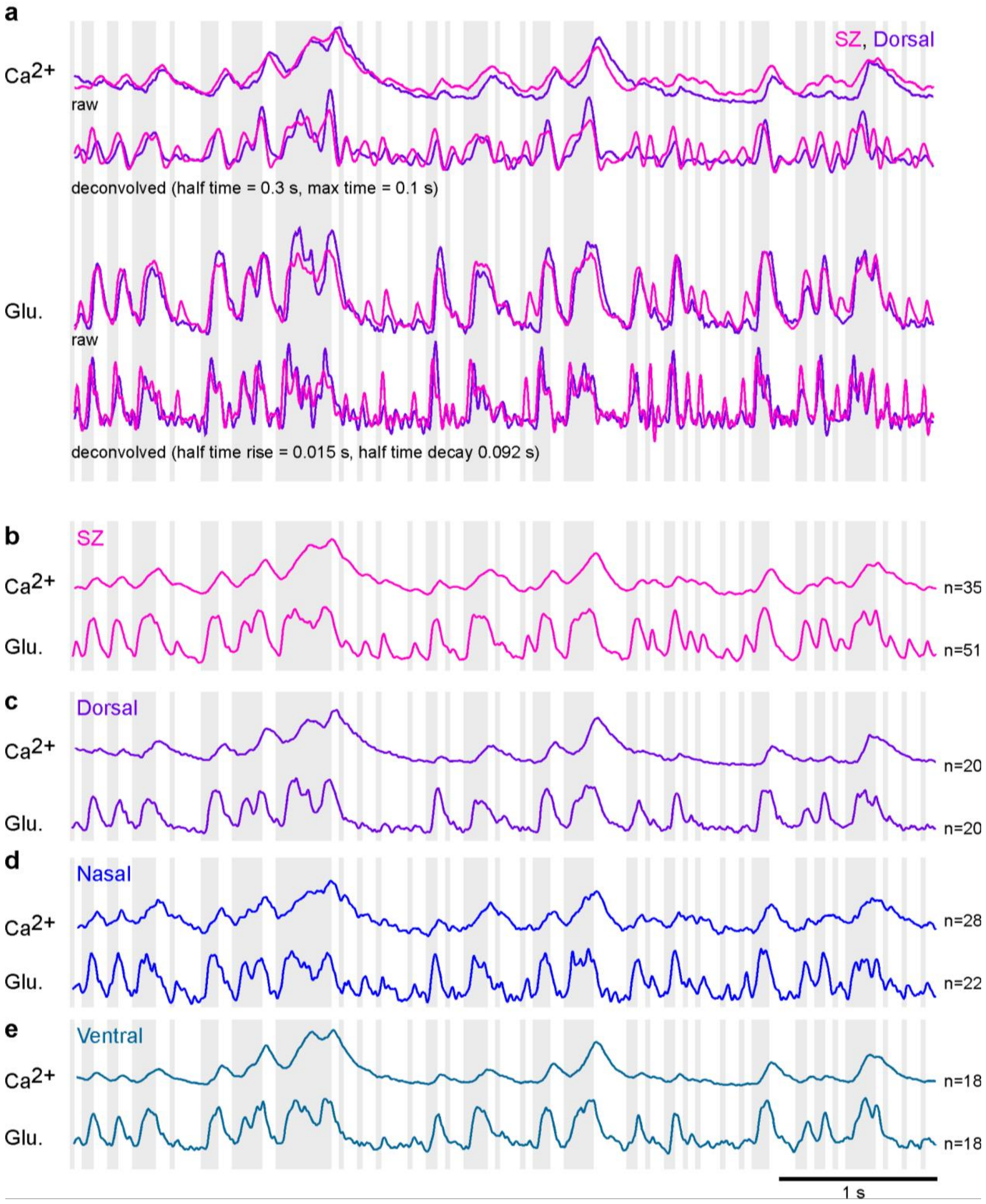


**Figure S3, related to Figure 8. Comparison of glutamate and calcium responses across retinal regions. a.** raw and deconvolved (Wiener deconvolution with calculated SNR using all recordings per zone. See also Method Section "Information rate") mean GCaMP6f[87] ($Ca^{2+}$, T = 0.3 s) and SFiGluSnFR[80,85] (glu., T = 0.092 s) responses from Fig. 8a to account for the kinetic differences between the sensors. The deconvolution does not strongly affect the differences between Dorsal and SZ UV-cones. **b-e.** Mean calcium and glutamate responses of UV-cones in the individual zones to the tetrachromatic noise stimulus. Background shading indicates UV-light and dark stimulus periods.

### SUPPLEMENTARY VIDEOS

**Supplementary Video S1, related to Figure 1. Detecting paramecia in UV and "yellow" wavebands**. Video of paramecia in naturalistic tank as viewed in a "yellow" channel that is approximately aligned with zebrafish M- and L-cones (left), and the same scene subsequently filmed in a zebrafish-approximate UV channel (right). The yellow channel provides spatial detail of the background and underside of the water, which masks paramecia swimming in the foreground. In contrast, the UV channel does not resolve the background clutter but instead brings out paramecia illuminated by the sun as bright dots in the upper water column. Videos recorded at 10 Hz and played back in real time (Methods).

**Supplementary Video S2, related to Figure 3. Imaging UV-cone synaptic calcium in vivo.** Calcium responses to bright- and dark-flashes in UV-cones from SZ (upper) and dorsal (D, bottom) as in Fig. 3b. The video is an average of 5 repeats of single trial raw movies that were cropped and aligned. The magenta bar indicates the timing of bright and dark flashes.

**Supplementary Video S3, related to Figure 4. A model of visual detectability of bright and dark moving objects.** Left, modelled UV-cone detector array (top) and bipolar cells (bottom) responding to a bright 2˚ target moving in a pseudorandom path at 100˚/s. The target is meant to mimic a paramecium. Right, as left, with target size increased to 5˚ and contrast inverted to dark. The target is meant to mimic a distant or small predator. In each case, the colour-scaling indicates relative activation of cones or bipolar cells scaled to the array's maximum. Note that the small light target is only readily detectable in the strike zone (top left in each array), while the predator is always detectable. Played back at real-time.

**Supplementary Video S4, related to Figure 5. Whole-eye imaging of light-driven UV-cone calcium levels.** UV-cone calcium responses to bright- and dark-flashes as in Fig. 5. The video is an average of 7 repeats of single trial raw movies that were cropped and aligned. The bars on the right indicate the timing of bright and dark flashes and the RGB background, which are all superimposed on a constant UV-background (not indicated).

**Supplementary Video S5, related to Figure 7. Imaging glutamate release from cones in vivo.** Video of mean glutamate responses over n=7 repetitions of the tetrachromatic binary noise stimulus as in Fig. 7. Green is SFiGluSnFR in HC and red is mCherry expression in UV-cones. The bars on the right indicate the timing of flashes of each LED.

**Supplementary Video S6, related to Figure 8. Glutamate release differences between SZ and dorsal.** Video of mean glutamate responses over n=4 repetitions of the tetrachromatic binary noise stimulus as in Fig. 7. Green is SFiGluSnFR in HC and red is mCherry expression in UV-cones. Circles indicate UV-cone terminals shown in the bottom as high-magnification. The bars on the right indicate the timing of flashes of each LED.
